# Rtn4a promotes exocytosis in mammalian cells while ER morphology does not necessarily affect exocytosis and translation

**DOI:** 10.1101/512129

**Authors:** Richik Nilay Mukherjee, Zhaojie Zhang, Daniel L. Levy

## Abstract

ER tubules and sheets conventionally correspond to smooth and rough ER, respectively. The ratio of ER tubules-to-sheets varies in different cell types and changes in response to cellular conditions, potentially impacting the functional output of the ER. To directly test if ER morphology impacts ER function, we increased the tubule-to-sheet ratio by Rtn4a overexpression and monitored effects on protein translation and trafficking. While expression levels of several cell surface and secreted proteins were unchanged, their exocytosis was increased. Rtn4a depletion reduced cell surface trafficking without affecting ER morphology, and increasing the tubule-to-sheet ratio by other means did not affect trafficking. These data suggest that Rtn4a enhances exocytosis independently of changes in ER morphology. We demonstrate that Rtn4a enhances ER-to-Golgi trafficking and co-localizes with COPII vesicles. We propose that Rtn4a promotes COPII vesicle formation by inducing membrane curvature. Taken together, we show that altering ER morphology does not necessarily affect protein synthesis or trafficking, but that Rtn4a specifically enhances exocytosis.

## INTRODUCTION

How organelle size and morphology affect organelle function is a fundamental question in cell biology. In this study, we investigated how morphology of the endoplasmic reticulum (ER) affects its functional output. The ER is an elaborate and dynamic membrane bound organelle that is continuous with the nuclear envelope. The ER network is composed of flat cisternae or sheets and arrays of highly curved tubules interconnected by three-way junctions (Shibata et al., 2006; Urra and Hetz, 2012; Goyal and Blackstone, 2013). Despite sharing a common luminal space, different ER domains are associated with distinct functions. Flat ER sheets correspond to rough ER (RER), accommodate large polyribosomes on their surface, and play major roles in protein translation, folding, and modification. Smooth ER (SER) is composed of ER tubules that generally exhibit low ribosome density because high membrane curvature disfavors binding of polyribosomal structures (Shibata et al., 2006; Goyal and Blackstone, 2013). Rather, ER tubules are involved in lipid synthesis, carbohydrate metabolism, calcium homeostasis, and interorganellar contacts (Park and Blackstone, 2010; Shibata et al., 2010; Goyal and Blackstone, 2013). Discrete specialized domains of the SER termed transitional ER (tER) or ER exit sites (ERES) represent sites of COPII coat assembly and vesicle budding (Hughes and Stephens, 2008). Vesicles packed with cargo proteins and lipids arise from ERES and traffic towards the Golgi apparatus through post-ER structures known as pre-Golgi intermediates (Lippincott-Schwartz et al., 2000; Hughes et al., 2009).

Multiple proteins contribute to the unique structures characteristic of different ER domains. ER tubules are curved by the Reticulon (Rtn) and DP1/REEP/Yop1p family of proteins, all of which contain two tandem hydrophobic hairpin segments, termed reticulon-homology domains (RHDs), thought to wedge into the outer leaflet of phospholipid bilayers to induce membrane curvature (Voeltz et al., 2006; Shibata et al., 2008; Goyal and Blackstone, 2013; Zhang and Hu, 2016). Although predominantly localized to ER tubules, reticulons also occupy the curved edges of ER sheets (Shibata et al., 2009; Shibata et al., 2010). There are four mammalian reticulon genes (*RTN1*, *RTN2*, *RTN3*, and *RTN4/Nogo*) that can give rise to alternatively spliced transcripts (Oertle and Schwab, 2003; Yang and Strittmatter, 2007). The C-terminal RHD is highly conserved among all reticulons, while their N-termini exhibit little or no sequence similarity (GrandPré et al., 2000). Rtn4a is the largest in the reticulon family and was originally identified as an inhibitor of neurite outgrowth and axonal regeneration in the central nervous system (Chen et al., 2000; GrandPré et al., 2000; Prinjha et al., 2000; Oertle and Schwab, 2003; Yang and Strittmatter, 2007; Zurek et al., 2011; Di Sano et al., 2012). While most reticulons, including Rtn4a and its shorter splice variants Rtn4b and Rtn4c, are enriched in the nervous system, they are also ubiquitously expressed in all tissues and localize to curved ER tubules and sheet edges (Chiurchiu et al., 2014; Ramo et al., 2016). The flat regions of ER sheets are supported by coiled-coil domain containing proteins, such as CLIMP-63, kinectin, and p-180. CLIMP-63 stabilizes a constant sheet width of 50-100 nm in mammals by forming intraluminal bridges, whereas p-180 and kinectin are thought to form rod-like structures on the surface of ER sheets to promote flatness (Klopfenstein et al., 2001; Voeltz and Prinz, 2007; Lin et al., 2012; Goyal and Blackstone, 2013).

Cells with specialized functions are enriched in specific ER morphologies. For example, pancreatic acinar cells and plasma cells, which produce and secrete large amounts of protein, are mostly populated with polyribosome studded ER sheets. In contrast, cells involved in carbohydrate metabolism (e.g. hepatocytes), steroid hormone synthesis (e.g. adrenal cortical cells), and Ca^+2^ signaling (e.g. muscle cells) are enriched in smooth tubular ER (Black, 1972; Shibata et al., 2006; Friedman and Voeltz, 2011; West et al., 2011; Goyal and Blackstone, 2013). The unfolded protein response and ER stress induced by excess fatty acids lead to expansion of ER sheets, which is energetically more favorable than ER tubule expansion and is thought to provide more space for protein folding (Schuck et al., 2009; Friedman and Voeltz, 2011; Wikstrom et al., 2013). Thus while correlations between ER morphology and function have been described in certain specialized cell types and in response to cellular conditions, the question remains whether ER morphology directly affects the functional output of the ER.

In our study, we manipulated the ER tubule-to-sheet ratio to test if this broadly impacts protein translation and trafficking, functional roles ascribed to the RER and SER, respectively. We overexpressed the tubule shaping protein Rtn4a/Nogo-A in HeLa cells to increase the ER tubule-to-sheet ratio, a previously validated approach to alter ER morphology (Voeltz et al., 2006; Puhka et al., 2007; Shibata et al., 2010; Romero-Brey and Bartenschlager, 2016). We examined a number of cell surface and secreted proteins, finding that while their total levels were unchanged upon Rtn4a overexpression, their trafficking through the secretory pathway was increased. We show that this effect on exocytosis is in fact not due to altered ER morphology, but is rather a specific function of Rtn4a. We also provide evidence that Rtn4a accelerates ER-to-Golgi trafficking by promoting COPII vesicle formation. Thus our data suggest that altering ER morphology does not necessarily influence protein translation and trafficking, but that Rtn4a has a specific function in promoting COPII-mediated exocytosis.

## RESULTS

### Rtn4a overexpression increases trafficking of cell surface proteins without changing their overall expression levels

HeLa cells were transiently transfected with plasmids expressing Rtn4a-GFP or GFP-NLS as a control, and overexpression of Rtn4a was confirmed by western blot and immunofluorescence (**Fig. S1, A-D**). Based on immunoblotting, Rtn4a levels were increased 8.8 ± 1.5 fold (average ± SD) compared to controls **(Fig. S1, A-B)**. This level of ectopic Rtn4a expression did not cause ER stress as evidenced by constant levels of the ER chaperones calnexin, ERp72, and GRP78 **(Fig. S1, E-H)**. To quantify Rtn4a-induced conversion of ER sheets into tubules, we stained cells for the ER sheet marker CLIMP63 **(Fig. S1, I)**. Rtn4a expression increased the relative proportion of tubular ER (**Fig. S1, C**) with a concomitant 2.6 ± 0.2 fold reduction in ER sheet volume **(Fig. S1, J)**. To assess how this change in ER morphology might affect protein synthesis and trafficking, we focused on two cell surface plasma membrane proteins whose transit through the secretory pathway has been well studied, Integrinβ1 and MHC class I/HLA-A (Akiyama et al., 1989; Gawantka et al., 1992; Jones et al., 1996; Sun et al., 2009; Rose et al., 2014). Total staining intensity for Integrinβ1 and HLA-A was unchanged upon Rtn4a overexpression, indicating that reducing ER sheet volume does not reduce synthesis of these two proteins (Fig. 1, E-H). Surprisingly, staining of non-permeabilized cells revealed greater cell surface levels of these membrane proteins in Rtn4a-transfected cells, by 1.4 ± 0.1 fold for Integrinβ1 and 3.2 ± 0.3 fold for HLA-A (Fig. 1, A-D). Rtn4a overexpression also increased the cell surface transport, but not total levels, of Integrinβ1 and HLA-A in MRC-5 cells, a non-cancerous lung fibroblast cell line **(Fig. S1, O-V)**. Thus in both a normal and cancer cell line, Rtn4a overexpression enhanced trafficking of Integrinβ1 and HLA-A to the cell surface without affecting their overall levels.

**Figure 1.**
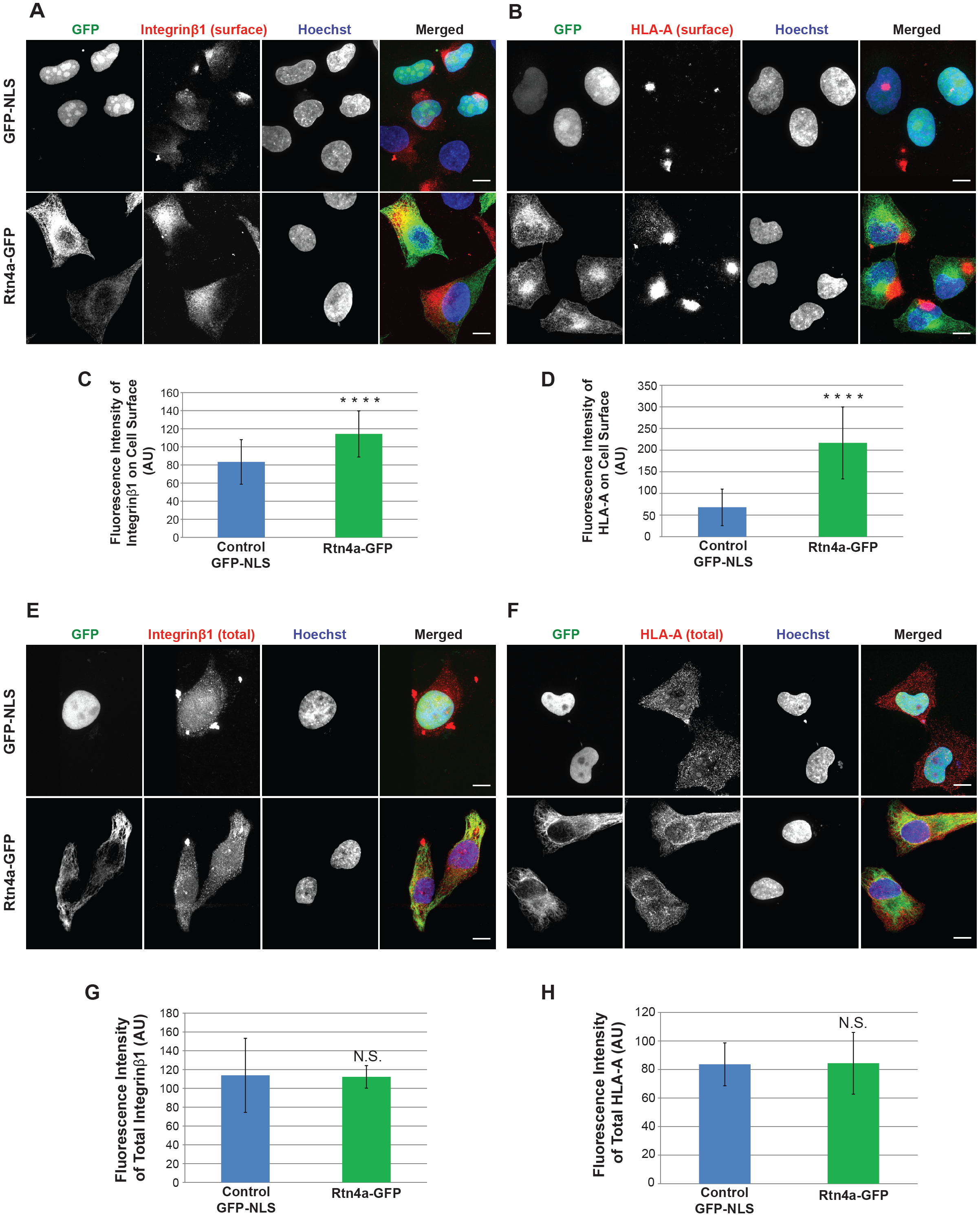
Rtn4a overexpression increases cell surface localization of Integrinβ1 and HLA-A, without changing their total cellular levels. HeLa cells were transiently transfected with plasmids expressing GFP-NLS as a control or Rtn4a-GFP (green). **(A-B)** Non-permeabilized cells were stained for surface-localized Integrinβ1 (**A**, red) or HLA-A (**B**, red) and DNA (blue). **(C)** Integrinβ1 surface fluorescence staining intensity was quantified for 39-52 cells per condition. **(D)** HLA-A surface fluorescence staining intensity was quantified for 59-69 cells per condition. **(E-F)** Permeabilized cells were stained for total Integrinβ1 (**E**, red) or HLA-A (**F**, red) and DNA (blue). **(G)** Total Integrinβ1 fluorescence staining intensity was quantified for 39-49 cells per condition. **(H)** Total HLA-A fluorescence staining intensity was quantified for 48-52 cells per condition. Scale bars are 10 µm and images are maximum intensity projections of confocal z-stacks. Error bars represent standard deviation. **** p≤0.0001; NS not significant.

We reasoned that enhanced trafficking of Integrinβ1 and HLA-A through the secretory pathway might lead to faster and overall increased maturation of these proteins. While immature N-glycoproteins in the ER lumen and cis-Golgi are sensitive to cleavage by endogycosidase H (EndoH), mature glycoproteins become EndoH-resistant after terminal mannose removal by mannosidase II in the medial Golgi. Whole cell lysates from control and Rtn4a-GFP transfected cells were treated with EndoH or buffer alone and immunoblotted for Integrinβ1 and HLA-A, allowing for quantification of mature, EndoH-resistant protein levels (Fig. 2, A and C). While total levels of Integrinβ1 and HLA-A were unaffected by Rtn4a overexpression, consistent with our immunofluorescence results, mature forms of Integrinβ1 and HLA-A were increased by 2.8 ± 0.8 fold and 1.8 ± 0.4 fold, respectively (Fig. 2, B and D). To assess all cell surface glycoproteins, non-permeabilized cells were stained with concanavalin A (ConA) which detects immature, high mannose N-glycans. Rtn4a overexpression decreased cell surface ConA-staining intensity by 1.2 ± 0.03 fold without affecting total ConA-staining levels **(Fig. S1, K-N)**, suggesting an increased presence of mature glycoproteins on the cell surface. Because the levels of key Golgi glycosyltransferases were unchanged upon Rtn4a overexpression (data not shown), Rtn4a does not appear to enhance protein maturation by modulating the protein glycosylation machinery. Instead, we propose that Rtn4a promotes exocytic trafficking, thereby leading to accelerated Golgi transport, N-glycosylation, and maturation. Taken together, these data show that Rtn4a overexpression, which converts ER sheets to tubules, promotes trafficking of two membrane glycoproteins to the cell surface without influencing their overall expression levels.

**Figure 2.**
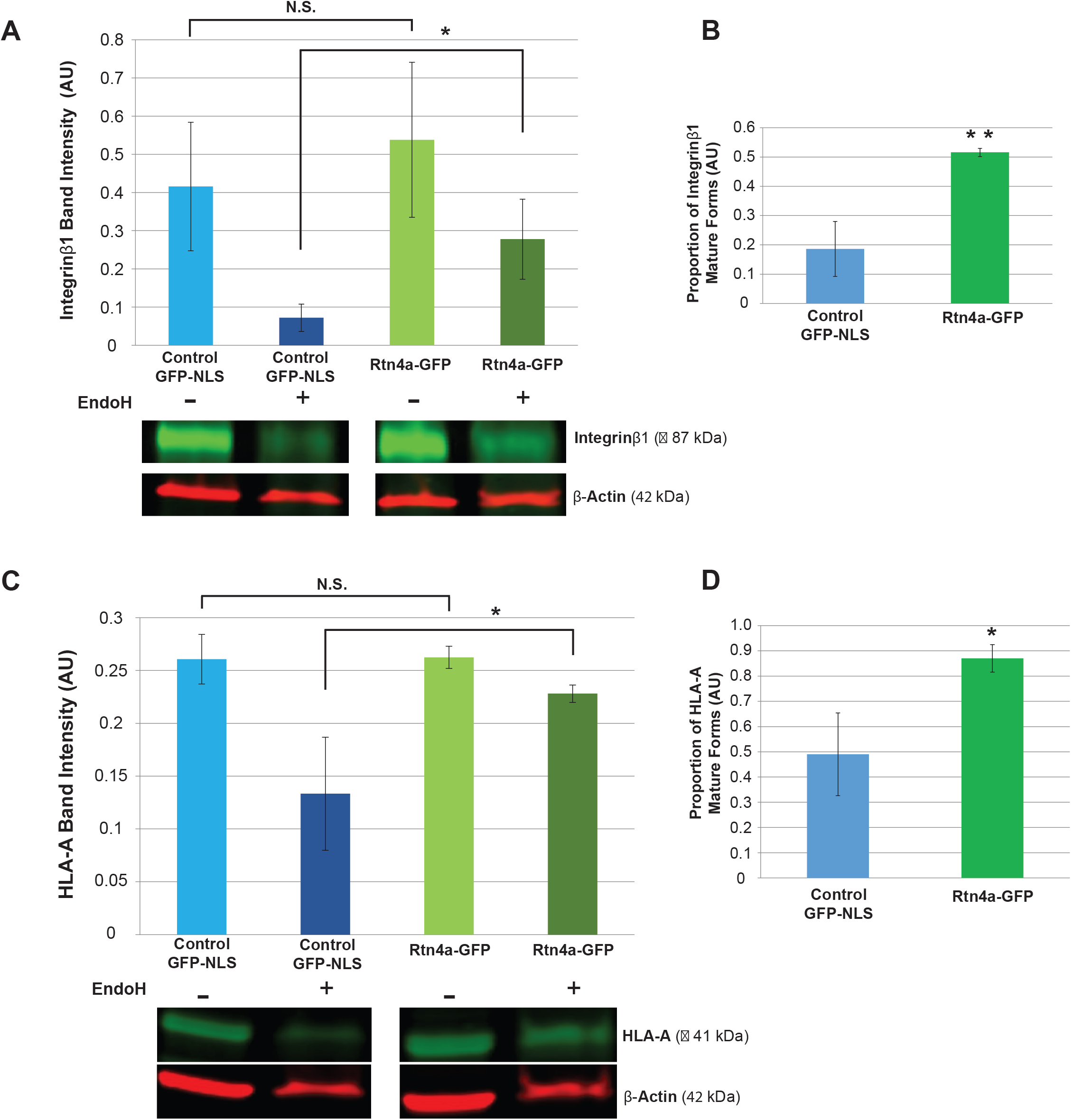
Rtn4a overexpression increases the proportion of mature forms of Integrinβ1 and HLA-A. HeLa cells were transiently transfected with plasmids expressing GFP-NLS as a control or Rtn4a-GFP. Whole cell lysates were subjected to gel electrophoresis, with or without prior endoglycosidase H digestion (EndoH), and immunoblotted for Integrinβ1, HLA-A, and β-Actin. **(A)** Integrinβ1 band intensities were quantified and normalized to β-Actin. **(B)** The proportion of Integrinβ1 mature forms was calculated by dividing the normalized EndoH-treated band intensity by the normalized untreated band intensity. **(C)** HLA-A band intensities were quantified and normalized to β-Actin. **(D)** The proportion of HLA-A mature forms was calculated by dividing the normalized EndoH-treated band intensity by the normalized untreated band intensity. Quantifications were performed from three independent experiments. Error bars represent standard deviation. ** p≤0.01; * p≤0.05; NS not significant.

### Rtn4a promotes trafficking of cell surface proteins independently of effects on ER morphology

While Rtn4a overexpression did not induce ER stress and increased trafficking of cell surface proteins, indicating that overexpression was not having dominant negative effects on cell function, we also wanted to test the consequences of Rtn4a depletion. HeLa cells transfected with siRNA against Rtn4 exhibited reduced Rtn4 expression (Fig. 3, A-B), with Rtn4a levels decreased by 4.3 ± 0.9 fold based on immunoblotting **(Fig. S2, A-C)**. Rtn4 knockdown reduced cell surface levels of Integrinβ1 and HLA-A by 1.2 ± 0.1 fold and 1.3 ± 0.1 fold, respectively (Fig. 3, C-F). Consistent with previous studies showing that Rtn1, 3, and 4 must be co-depleted to allow for conversion of ER tubules into sheets (Voeltz et al., 2006; Anderson and Hetzer, 2008; Christodoulou et al., 2016), we observed no change in ER sheet volume in Rtn4 knockdown cells (Fig. 3, G-H). These results suggest that Rtn4 influences protein trafficking independently of any effect on ER morphology.

**Figure 3.**
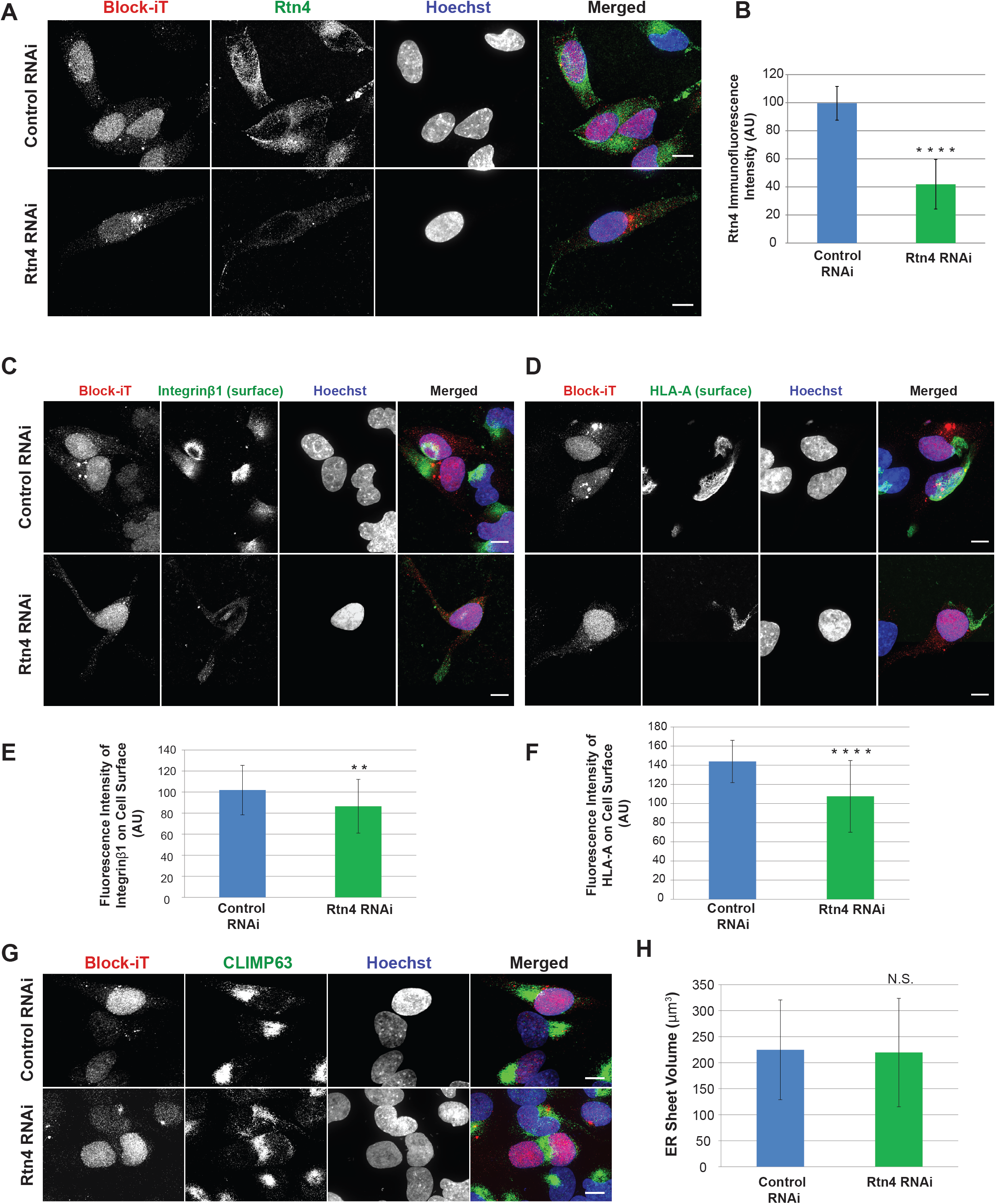
Knockdown of Rtn4 decreases cell surface localization of Integrinβ1 and HLA-A without affecting ER sheet volume. HeLa cells were transiently co-transfected with siRNA against Rtn4 and Block-iT fluorescent control siRNA or with Block-iT alone (red). **(A)** Cells were immunostained for Rtn4 (green) and DNA (blue). **(B)** Average fluorescence intensity of Rtn4 immunostaining was quantified for 30-36 cells per condition. **(C-D)** Non-permeabilized cells were stained for surface-localized Integrinβ1 (**C**, green) or HLA-A (**D**, green) and DNA (blue). **(E)** Integrinβ1 surface fluorescence staining intensity was quantified for 39-51 cells per condition. **(F)** HLA-A surface fluorescence staining intensity was quantified for 34-51 cells per condition. **(G)** Cells were stained for ER sheet marker CLIMP63 (green) and DNA (blue). **(H)** Mean ER sheet volume based on CLIMP63 immunofluorescence was quantified from 3D reconstructed confocal z-stacks. 35-57 cells were quantified for each condition. Scale bars are 10 µm. Images are maximum intensity projections of confocal z-stacks. Error bars represent standard deviation. **** p≤0.0001; ** p≤0.01; NS not significant.

To test if altering ER morphology through Rtn4a-independent means would impact exocytosis, we overexpressed two other curvature stabilizing ER membrane proteins – REEP5 and Rtn4b. Overexpressing REEP5 decreased ER sheet volume 2.9 ± 0.04 fold **(Fig. S2, D-G)**, similar to the 2.6-fold reduction in ER sheet volume induced by Rtn4a overexpression **(Fig S1, I-J)**, but had no effect on the amount of surface localized Integrinβ1 and HLA-A **(Fig. S2, H-J)**. Consistent with these results, a 6.9 ± 1.8 fold increase in Rtn4b expression led to a 2.3 ± 0.2 fold reduction in ER sheet volume **(Fig. S2, K-N)** without changing the levels of Integrinβ1 and HLA-A on the cell surface **(Fig. S2, O-Q)**. Collectively, these data show that shifting ER morphology from sheets to tubules is not sufficient to enhance trafficking of membrane proteins to the cell surface, implying Rtn4a has a unique ER morphology-independent function in this process.

### Overexpression of Rtn4a increases the secretion of soluble proteins

To test if Rtn4a promotes exocytosis of soluble proteins in addition to cell surface membrane proteins, we examined the secreted and intracellular levels of fibulin-5 (FBLN5) and thrombospondin-1 (TSP1). FBLN5 and TSP1 are both extracellular matrix components ubiquitously expressed and secreted by many cells types (Lahav, 1993; Bornstein, 1995; Crawford et al., 1998; Albig and Schiemann, 2005). Rtn4a overexpression increased the amount of FBLN5 and TSP1 secreted into the media (Fig. 4, A and D), without affecting their overall expression levels (Fig. 4, C and F). While the intracellular FBLN5 concentration was unchanged upon Rtn4a overexpression (Fig. 4 B), the intracellular TSP1 concentration was reduced (Fig. 4 E), suggesting TSP1 might be trafficked faster than FBLN5. These data show that Rtn4a promotes exocytosis of soluble secreted proteins in addition to membrane-bound proteins.

**Figure 4.**
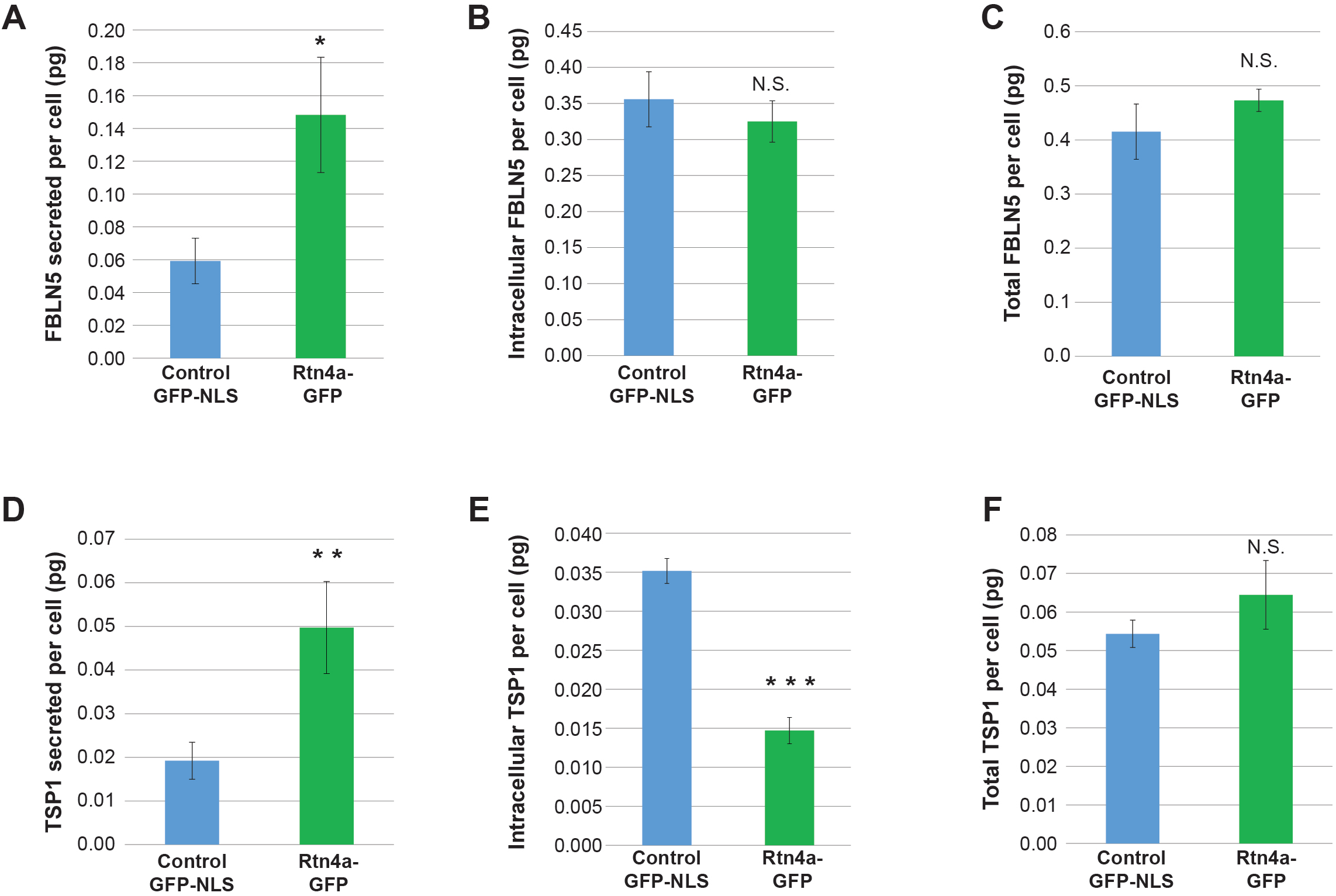
Overexpression of Rtn4a increases the secretion of endogenous FBLN5 and TSP1. HeLa cells were transiently nucleofected with plasmids expressing GFP-NLS as a control or Rtn4a-GFP. Media was collected and whole cell lysates were prepared 12 hours post transfection. Samples were subjected to sandwich ELISA. **(A,D)** Amounts of secreted FBLN5 (A) and TSP1 (D) present in the media were quantified by ELISA and normalized to the number of live cells. **(B,E)** Amounts of intracellular FBLN5 (B) and TSP1 (E) present in whole cell lysates were quantified by ELISA and normalized to the number of live cells. **(C,F)** Total amounts of FBLN5 (C) and TSP1 (F) were calculated by summing the secreted and intracellular amounts of each protein per cell. Data are presented from three independent experiments. Error bars represent standard deviation. *** p≤0.001; ** p≤0.01; * p≤0.05; NS not significant.

### Rtn4a accelerates ER-to-Golgi trafficking

To begin to address how Rtn4a might promote protein trafficking to the cell surface, we used the RUSH system to monitor transport of a fluorescent cargo through the exocytic pathway. HeLa cells were transiently co-transfected with a LAMP1-RUSH construct encoding mCherry-LAMP1 and streptavidin-Ii (Boncompain et al., 2012), along with Rtn4a-GFP or GFP-NLS as a control. Prior to biotin addition, mCherry-LAMP1 is trapped in the ER (Fig. 5 A; first row of images). After biotin addition and LAMP1 release from the ER, we fixed cells at 15-minute intervals. While control cells exhibited compact LAMP1 localization in the perinuclear region 15 minutes after biotin addition, in Rtn4a-overexpressing cells LAMP1 was present in puncta dispersed throughout the cytoplasm (Fig. 5 A; second row of images). Similar LAMP1 cytoplasmic puncta were not observed in control cells until 45 minutes after biotin addition (Fig. 5 A; fourth row of images). These data suggest that LAMP1 exocytosis occurs more rapidly when Rtn4a is overexpressed.

**Figure 5.**
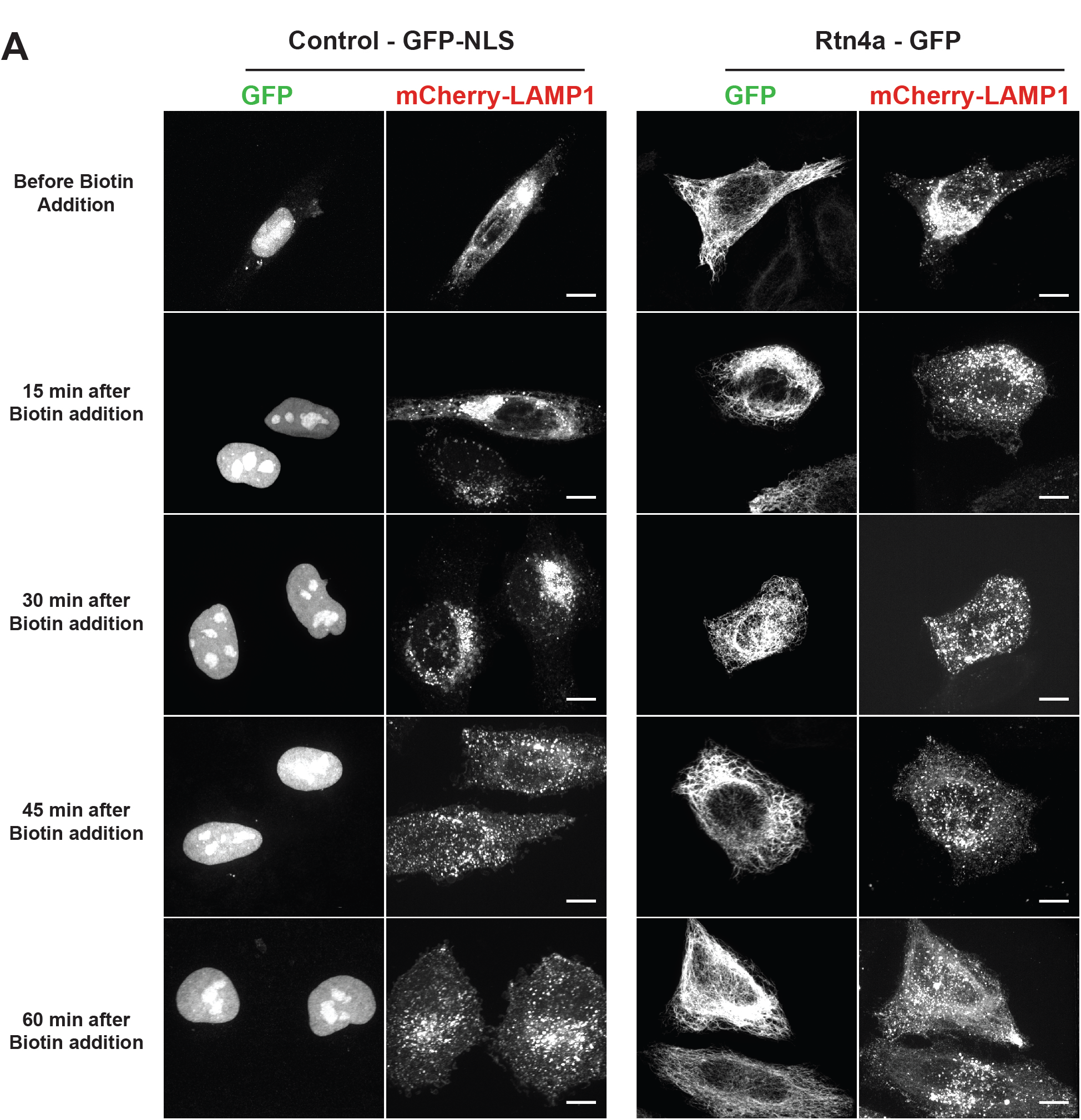
Overexpression of Rtn4a accelerates trafficking of ectopically expressed mCherry-LAMP1. HeLa cells were transiently co-transfected with plasmids expressing GFP-NLS as a control or Rtn4a-GFP (green) and the RUSH construct Str-Ii_LAMP1-SBP-mCherry (red). ER-trapped LAMP1 was released by addition of 40 µM D-Biotin to the growth media. **(A)** Cells fixed before biotin addition and at the indicated 15-minute intervals were imaged. Scale bars are 10 µm. Images are maximum intensity projections of confocal z-stacks.

To estimate when LAMP1 was entering the Golgi, cells were fixed at shorter time points after biotin addition and immunostained with the Golgi marker GRASP65 to assess co-localization of LAMP1 with the Golgi (**Fig. S3, A-D**). Just a few minutes after biotin addition, we observed significantly greater co-localization of LAMP1 with the Golgi in cells overexpressing Rtn4a compared to controls **(Fig. S3 E)**, suggesting accelerated ER-to-Golgi transport. This prompted us to track ER-to-Golgi transport of LAMP1 by live cell imaging (Fig. 6). We monitored the accumulation of mCherry-LAMP1 signal in the presumptive Golgi as an increase in perinuclear fluorescence relative to peripheral ER fluorescence. Perinuclear LAMP1 fluorescence peaked 6 minutes after biotin addition in Rtn4a-overexpressing cells (Fig. 6, D-F and H; **Video S1B**), 9 minutes earlier than in control cells (Fig. 6, A-C and G; **Video S1A**). It appeared that the Golgi was more disperse in Rtn4a-transfected cells (Fig. 6 E; 6 minutes after biotin addition) compared to control cells (Fig. 6 B; 15 minutes after biotin addition), which we confirmed by staining for GRASP65 and the medial Golgi marker ManII, observing a ~2-fold decrease in Golgi circularity and ~2-3 fold increase in Golgi volume upon Rtn4a overexpression (Fig. 7, **S3 F**). Golgi morphology was also altered in REEP5- and Rtn4b-overexpressing cells in which cell surface trafficking was unaffected (**Fig. S3, G-I**), suggesting that Golgi fragmentation and enlargement are not sufficient to increase exocytosis and that Rtn4a likely influences anterograde trafficking through another mechanism. Taken together, these data suggest that Rtn4a overexpression accelerates ER-to-Golgi transport, which in turn increases trafficking of proteins to the cell surface.

**Figure 6.**
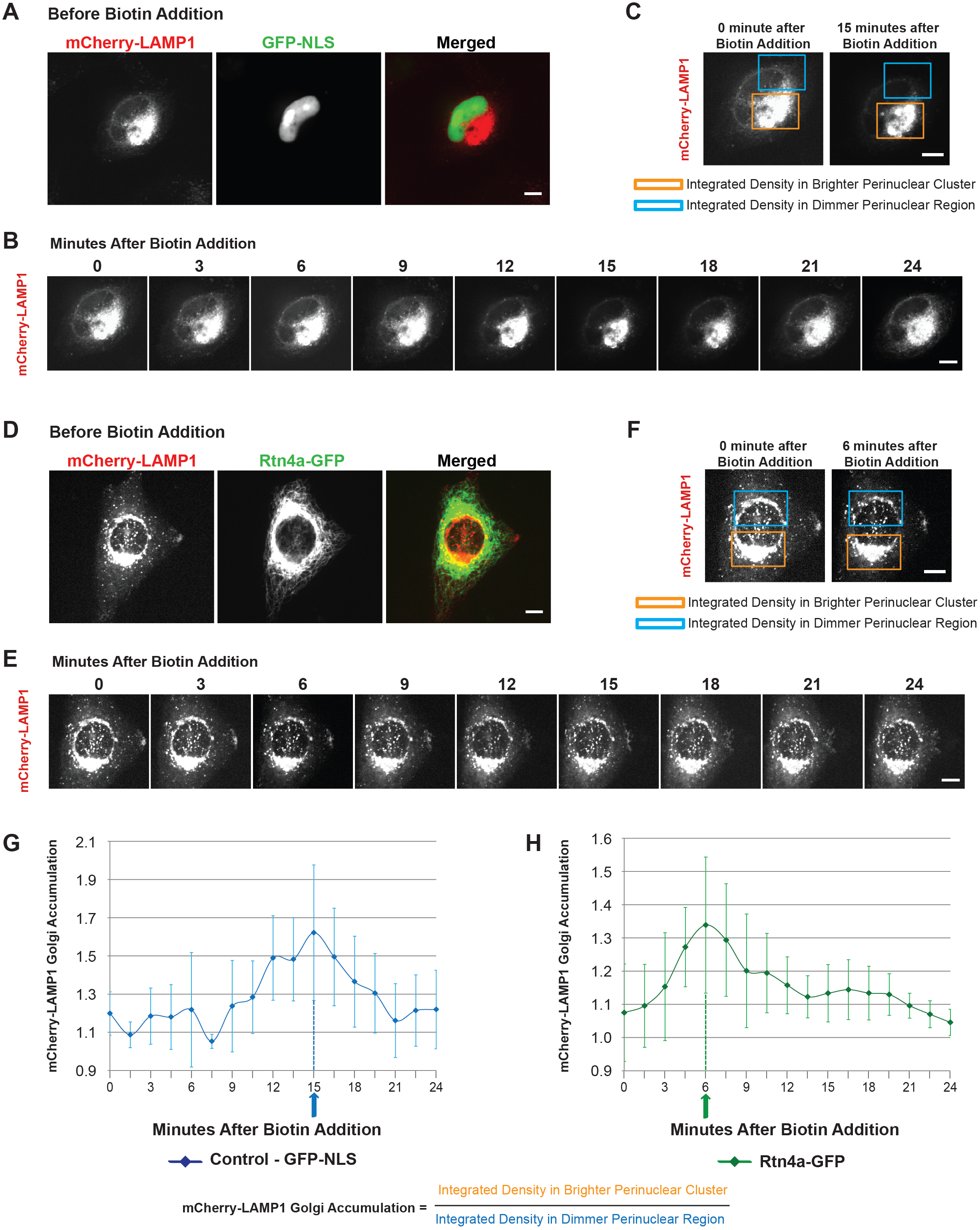
Rtn4a overexpression accelerates trafficking of LAMP1 from the ER to the Golgi. HeLa cells were transiently co-transfected with plasmids expressing GFP-NLS as a control or Rtn4a-GFP (green) and the RUSH construct Str-Ii_LAMP1-SBP-mCherry (red). ER-trapped LAMP1 was released by addition of 40 µM D-Biotin to the growth media. Live cell imaging was performed at 90-second intervals. **(A,D)** Representative images prior to biotin addition. **(B,E)** Representative images at 3-minute intervals after biotin addition. **(C,F)** In control cells (C) and Rtn4a-GFP transfected cells (F), integrated density of mCherry-LAMP1 fluorescence was measured for the indicated brighter perinuclear clusters and dimmer perinuclear regions. **(G-H)** To estimate accumulation of LAMP1 in the Golgi over time, the integrated density of the brighter perinuclear LAMP1 signal was divided by the integrated density of the dimmer perinuclear LAMP1 signal at each time point. The peaks indicated by arrows at 15 minutes for the control and 6 minutes for Rtn4a overexpression represent the highest relative accumulation of LAMP1 in the Golgi. 4-5 cells were quantified per condition based on live time-lapse imaging of single z-planes. Scale bars are 10 µm. Error bars represent standard deviation.

**Figure 7.**
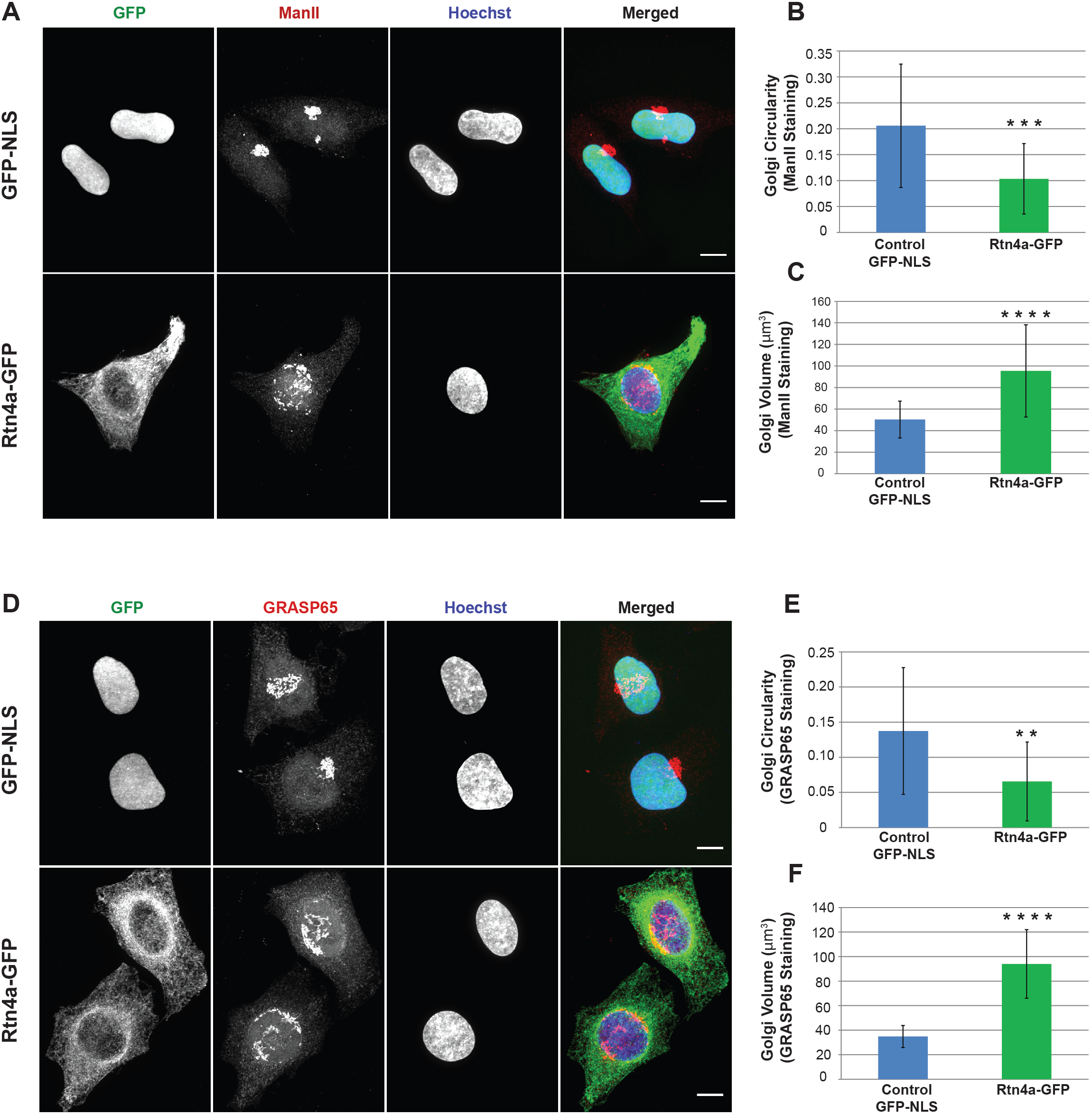
Rtn4a overexpression causes Golgi fragmentation and enlargement. HeLa cells were transiently transfected with plasmids expressing GFP-NLS as a control or Rtn4a-GFP (green). **(A)** Cells were stained for medial Golgi marker ManII (red) and DNA (blue). **(B-C)** Golgi circularity (B) and volume (C) were quantified based on ManII staining for 26-29 cells per condition. Volume was quantified from 3D reconstructed confocal z-stacks. **(D)** Cells were stained for Golgi marker GRASP65 (red) and DNA (blue). **(E-F)** Golgi circularity (E) and volume (F) were quantified based on GRASP65 staining for 38-41 cells per condition. Volume was quantified from 3D reconstructed confocal z-stacks. Scale bars are 10 µm and images are maximum intensity projections of confocal z-stacks. Error bars represent standard deviation. **** p<0.0001; *** p<0.001; ** p<0.01.

### Overexpressed Rtn4a increases Sec31A staining area and co-immunoprecipitates with Sec31A vesicles

To begin to understand how Rtn4a might influence ER-to-Golgi vesicular trafficking, we immunostained cells with the ER exit site (ERES) marker Sec31A, a key component of the outer COPII coat that facilitates budding of anterograde vesicles out of the ER (Lippincott-Schwartz et al., 2000; Fromme and Schekman, 2005; Melero et al., 2018). While predominantly perinuclear in control cells, Sec31A appeared more scattered throughout the cytoplasm of Rtn4a-overexpressing cells and the Sec31A staining area was 1.4 ± 0.2 fold greater (Fig. 8, A-B). Intriguingly, Sec31A puncta frequently co-aligned with cortical ER tubules labeled with Rtn4a-GFP (Fig. 8, A and C). Furthermore, in Sec31A co-immunoprecipitation experiments, overexpressed Rtn4a co-precipitated with immuno-isolated intact COPII vesicles, as did a small fraction of endogenous Rtn4a in control-transfected cells (Fig. 8, D-E). These data suggest that Rtn4a may promote budding of COPII vesicles, perhaps by inducing membrane curvature at ERES. Consistent with Rtn4a promoting anterograde trafficking, Rtn4a overexpression also led to more disperse staining of ERGIC53 **(Fig. S4, A-D)**, a receptor for glycoprotein transport from ERES and a marker of transient pre-Golgi structures found in the ER-Golgi intermediate compartment (ERGIC) (Appenzeller et al., 1999; Lippincott-Schwartz et al., 2000; Fromme and Schekman, 2005). Moreover, Rtn4a overexpression scattered the distribution of COPI-coated retrograde vesicles **(Fig. S4, E-G**), suggesting that Golgi-to-ER retrograde traffic may also increase upon Rtn4a overexpression, possibly balancing enhanced forward transport (Lippincott-Schwartz et al., 2000).

**Figure 8.**
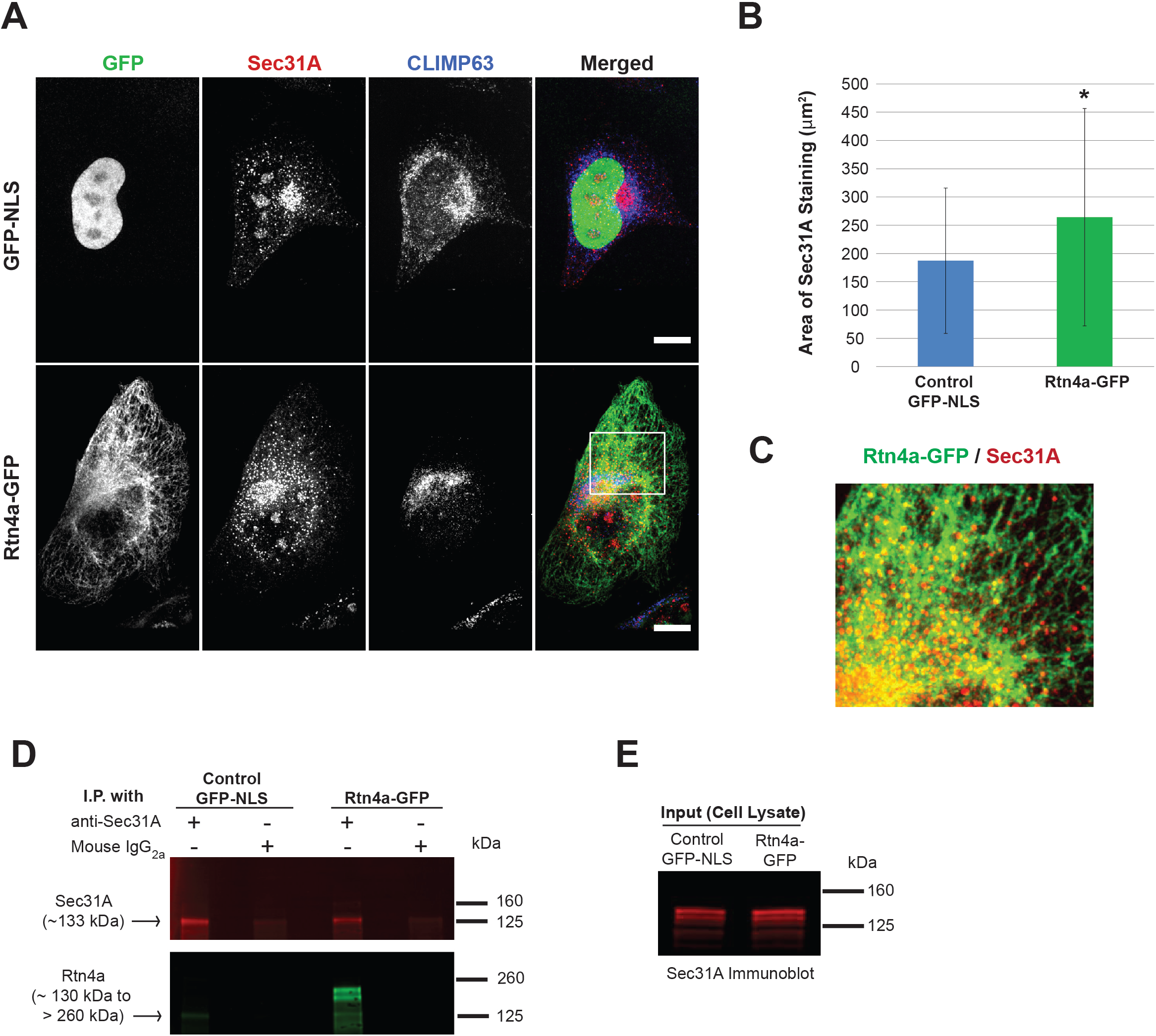
Overexpressed Rtn4a increases Sec31A staining area and co-immunoprecipitates with Sec31A vesicles. HeLa cells were transiently transfected with plasmids expressing GFP-NLS as a control or Rtn4a-GFP (green). **(A)** Cells were immunostained for COPII coat marker Sec31A (red) and ER sheet marker CLIMP63 (blue). **(B)** Total area of Sec31A staining was quantified for 52-68 cells per condition. **(C)** A magnified image of the white box in (A) showing the Rtn4a-GFP/Sec31A merge in an Rtn4a overexpressing cell. Sec31A shows close alignment with ER tubules. **(D-E)** Detergent-free cell homogenates with intact membranes from control and Rtn4a-GFP transfected cells were subjected to immunoprecipitation using an anti-Sec31A antibody or mouse IgG_2a_. In (D), immunoprecipitated samples were immunoblotted for Sec31A (top) and Rtn4a (bottom). In (E), whole cell lysates were immunoblotted for Sec31A. Scale bars are 10 µm. Images are maximum intensity projections of confocal z-stacks. Error bars represent standard deviation. * p≤0.05.

### Overexpressed Rtn4a closely associates with Sec31A-containing vesicles and tubules

To test if overexpressed Rtn4a might promote recruitment of COPII coat components, we performed transmission electron microscopy on cells immuno-labeled for Rtn4a and Sec31A with 6 nm and 15 nm gold particles, respectively. Rtn4a and Sec31A were more frequently observed within the same tubular or vesicular compartments in Rtn4a-overpressing cells compared to control cells (Fig. 9, A-B). Because ER tubules are 50-100 nm in diameter (Bernales et al., 2006; Shibata et al., 2006) and COPII-coated vesicles are generally 60-70 nm wide (Fromme and Schekman, 2005), we reasoned that in order for Rtn4a and Sec31A to co-localize in the same membrane compartment, 6 and 15 nm gold particles should be at most 100 nm apart from each other. We therefore quantified the number of 6 nm particles within 100 nm of each 15 nm particle, which was 3.9 ± 0.3 fold greater in Rtn4a-overexpressing cells compared to control cells (Fig. 9 C). Because this difference might be due to Rtn4a overexpression itself, we also quantified the density of 6 nm particles in regions devoid of Sec31A, finding that the Rtn4a particle density was 1.6 ± 0.1 fold higher close to Sec31A as opposed to far from Sec31A in Rtn4a-overexpressing cells (Fig. 9 C). These data suggest that COPII coats are preferentially recruited to Rtn4a-containing membranes. Consistent with this idea, we also measured the distance between each 15 nm (Sec31A) particle and the nearest 6 nm (Rtn4a) particle, observing a 1.8 ± 0.1 fold decrease in this distance upon overexpression of Rtn4a (Fig. 9 D).

**Figure 9.**
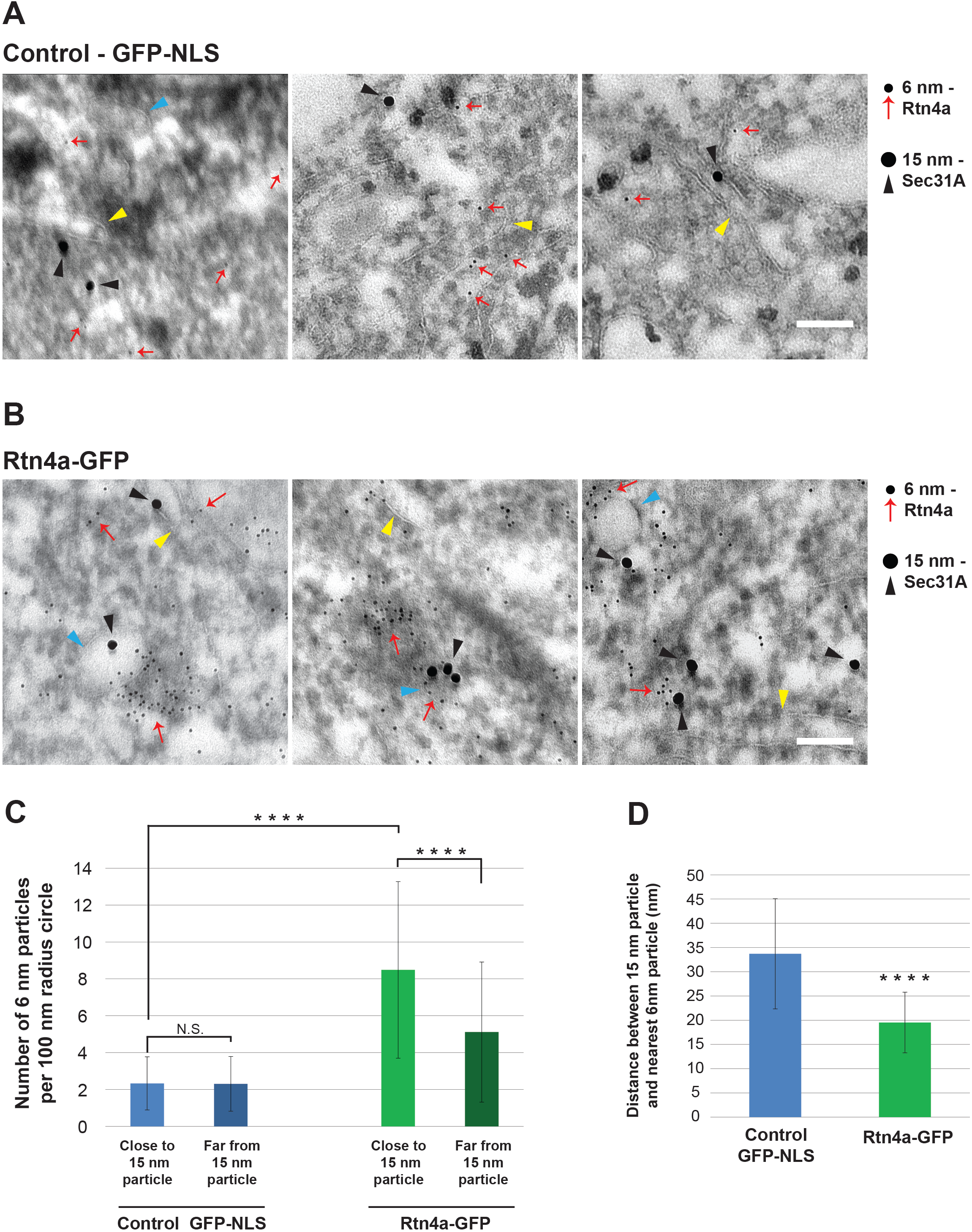
Overexpressed Rtn4a closely associates with Sec31A-containing vesicles and tubules. HeLa cells were transiently transfected with plasmids expressing GFP-NLS as a control or Rtn4a-GFP. Cells were processed for TEM, and immuno-gold labeling was performed to label Rtn4 with 6 nm gold particles and Sec31A with 15 nm gold particles. **(A-B)** Representative immuno-TEM micrographs of cells showing membrane bound vesicles (turquoise arrowheads) and tubules (yellow arrowheads). Red arrows denote single or multiple 6 nm gold particles corresponding to Rtn4a. 15 nm gold particles marked by black arrowheads represent Sec31A. **(C)** To quantify the density of 6 nm particles close to a 15 nm particle, the number of 6 nm particles was counted within a 100 nm radius circle around a 15 nm particle (85 – 127 15 nm particles per condition). To quantify the density of 6 nm particles in regions devoid of 15 nm particles, the number of 6 nm particles was counted within randomly selected 100 nm radius circles (255 - 286 randomly selected 100 nm radius circles per condition). **(D)** The distances between a 15 nm particle and the nearest 6 nm particle were quantified for 85-129 15 nm particles per condition. 7-10 50 nm ultra-thin sections were analyzed per condition. Scale bars are 100 nm. Error bars represent standard deviation. **** p≤0.0001; NS not significant.

We also observed co-localization of Sec31A and Rtn4a in membrane structures that appeared morphologically distinct from canonical membrane vesicles **(Fig. S5, A-B)**. These membrane structures protruded from enclosed membrane compartments and consisted of narrow tubular necks terminating in wider vesicular heads. Rtn4a was frequently observed in these structures and Sec31A tended to localize along the necks **(Fig. S5, A-B)**. Perhaps Rtn4a can promote an alternative mode of vesicle budding at ERES whereby the COPII machinery pinches off a tubular vesicle instead of coating the vesicle surface. We also quantified a 1.2 ± 0.03 fold decrease in ER tubule width upon Rtn4a overexpression **(Fig. S5 C)**, consistent with the idea that Rtn4a-induced membrane constriction and curvature might promote recruitment of the COPII vesicle budding machinery.

## DISCUSSION

We report that inducing a more tubular ER does not significantly influence protein synthesis levels in both HeLa cells and a normal lung fibroblast cell line. ER sheets are synonymous with rough ER, bound by translating polyribosomes and associated with protein biosynthesis, folding, and post-translational modification (Shibata et al., 2006). Conversely, smooth ER tubules are largely devoid of polyribosomes and are instead specialized for calcium signaling and lipid metabolism. Why might reducing ER sheet volume have no effect on protein synthesis levels? One possibility is that active polyribosomes redistribute to the tubular ER in Rtn4a-overexpressing cells. However, our TEM images provided no evidence of increased polyribosomes on ER tubules (data not shown), consistent with the observation that ER tubules are only populated by non-translating single ribosomes and small polyribosomes (Shibata et al., 2006; Shibata et al., 2010), likely because the high curvature of the SER prevents recruitment of large, spiral polysomal structures. Rather we propose that the biosynthetic capacity of rough ER sheets is sufficiently high so that a ~2.6-fold reduction in ER sheet volume has no effect on overall protein levels. Consistent with this idea, our TEM images showed no significant change in the density of spiral polyribosomes on the RER upon Rtn4a overexpression (data not shown). Furthermore, converting ER tubules into sheets by overexpression of CLIMP63 did not increase synthesis of Integrinβ1 and HLA-A (data not shown), again suggesting that the relative luminal volume of the RER does not directly impact the output of the translational machinery. Thus while there are specialized cell types that are enriched in rough ER sheets or tubular smooth ER (Friedman and Voeltz, 2011; Goyal and Blackstone, 2013), altering ER morphology in cells with a more traditional ER tubule-to-sheet ratio has little effect on overall protein synthesis levels.

While Rtn4a overexpression did not alter levels of protein synthesis, we did observe an increase in ER-to-Golgi trafficking and exocytosis of both membrane-bound and soluble proteins. This effect was likely not due to a change in ER morphology, as cell surface trafficking was unaffected by overexpression of two other ER tubule-promoting proteins, Rtn4b and REEP5, and Rtn4a knock down reduced trafficking without affecting ER morphology. Thus Rtn4a appears to have a specific function in enhancing anterograde transport. If ER morphology has little effect on protein translation and trafficking, how might Rtn4a specifically promote exocytosis? Rtn4a is known to induce membrane curvature (Voeltz et al., 2006; Shibata et al., 2008; Zurek et al., 2011), and COPII vesicle budding preferentially occurs at sites of high membrane curvature (Wang et al., 2017; Melero et al., 2018). As our immunofluorescence, co-immunoprecipitation, and immuno-TEM experiments all suggested co-localization of Rtn4a and Sec31A, we propose that membrane curvature induced by Rtn4a promotes recruitment of the COPII coat machinery, thereby facilitating vesicle budding from ERES and enhancing overall exocytosis. REEP5 and Rtn4b overexpression did not alter protein transport to the cell surface, indicating that membrane curvature by itself cannot enhance trafficking and that Rtn4a must contain a unique structural element that confers this ability. The splice isoform Rtn4b lacks a unique N-terminal domain present in Rtn4a, suggesting that this region may be involved in promoting trafficking. It is worth noting that this same N-terminal region is also associated with Rtn4a’s inhibitory effect on neurite outgrowth (GrandPré et al., 2000; Oertle and Schwab, 2003; Yan et al., 2006; Yang and Strittmatter, 2007), suggesting a potential link between these two functions. While C-terminal transmembrane domains in Rtn4 are known to induce membrane curvature, there is evidence that other domains of Rtn4a may also be important in shaping the tubular ER (Zurek et al., 2011). Perhaps unique biophysical properties of Rtn4a-induced membrane curvature contribute to enhanced exocytosis. Future studies will elucidate the structural elements of Rtn4a that enhance vesicular transport.

Our results with Rtn4a add to a growing literature on the role of reticulons in protein trafficking. Overexpression of RTN1C in rat PC12 adrenal tumor cells increased exocytosis of human growth hormone, mediated by interactions with the SNARE proteins syntaxin-1, −7, −13, and VAMP2 (Steiner et al., 2004; Di Sano et al., 2012). RTN3 has been shown to play a role in retrograde protein transport from the cis-Golgi and/or ERGIC to the ER. Notably, overexpression of RTN3 in HeLa cells delayed trafficking of VSVG from the ER to the cell surface and restricted ERGIC53 staining to the perinuclear region (Wakana et al., 2005). Combining these results with our own, perhaps Rtn3 and Rtn4a have antagonizing effects on the early secretory pathway. Previous studies support a role for Rtn4a in neuronal protein secretion. Rtn4a mRNA is highly expressed in the supraoptic nucleus and paraventricular nucleus of the rat hypothalamus, both regions being highly active for neuroendocrine secretion (Hasegawa et al., 2005), and RTN4A knockdown decreased dopamine release in rat PC12 cells (Xiong et al., 2008). Thus, while Rtn4a/Nogo-A has been implicated in protein secretion in the nervous system, our results show that Rtn4a plays a more general role in exocytosis, providing a potential mechanism that involves recruitment of the COPII vesicle budding machinery.

Finally, Rtn4a’s capacity to enhance exocytosis might be relevant in cancer and neurodegenerative disease, where pathogenesis is often driven by abnormal secretion of growth factors, proteases, and neuropeptides (Daughaday and Deuel, 1991; Lynch and Mobley, 2000; Yan et al., 2006; Lodish et al., 2008). For example, Rtn4a protein is upregulated in malignant brain tumor (glioma) cells (Björling et al., 2008), concomitant with increased secretion of cathepsin B (McCormick, 1993), VEGF (Jensen et al., 2006), and chemokines (Jordan et al., 2008). Amyotrophic Lateral Sclerosis (ALS) is associated with motor neuron degeneration and skeletal muscle paralysis. High levels of Rtn4a were measured in the muscles of ALS patients as well as in mouse models of ALS. Knocking down RTN4A in ALS mice delayed disease progression and extended survival (Dupuis et al., 2002; Bruijn et al., 2004; Jokic et al., 2006). Given that Rtn4a is an inhibitor of neurite outgrowth and axonal regeneration, processes regulated by vesicular trafficking (Tojima and Kamiguchi, 2015), perhaps Rtn4a contributes to ALS by influencing exocytosis (Yang and Strittmatter, 2007). The pathogenesis of Parkinson’s disease has also been linked to Rtn4a (More et al., 2013; Schawkat et al., 2015), with a possible mechanism being increased secretion of the inflammatory cytokines TNFα and IL-6 (Zhong et al., 2015). Future research will examine the potential disease links between Rtn4a and protein trafficking.

## MATERIALS AND METHODS

### Plasmids and siRNA

The Rtn4a-GFP (pAcGFP1-N1 Rtn4a) (Shibata et al., 2008) and mCherry-REEP5 (pmCherry-C2 REEP5) (Schlaitz et al., 2013) mammalian expression plasmids were generous gifts from Gia Voeltz (University of Colorado, Boulder) and Anne Schlaitz (Zentrum für Molekulare Biologie der Universität Heidelberg), respectively. The RUSH construct Str-Ii_LAMP1-SBP-mCherry was a gift from Franck Perez (Boncompain et al., 2012). The GFP-Rtn4b expression plasmid (pDL34) was described previously (Jevtić and Levy, 2015). The control GFP-NLS plasmid was from Invitrogen (V821-20). To knock down expression of RTN4, we used a DsiRNA (IDT): sense 5’-CUGGAAUCUGAAGUUGCUAUAUCUG-3’, and antisense 5’-CAGAUAUAGCAACUUCAGAUUCCAG-3’. This DsiRNA sequence is based on a previously described siRNA against RTN4 (Anderson and Hetzer, 2008).

### Mammalian Tissue Culture and Transfections

HeLa cells were obtained from ATCC and MRC-5 normal human lung fibroblast cells were a gift from Jason Gigley (University of Wyoming). Cells were verified to be mycoplasma-free (ThermoFisher Scientific #M7006). Both cell lines were cultured in Eagle’s minimum essential medium (EMEM), supplemented with 10% v/v fetal bovine serum (FBS) and 50 IU/ml of penicillin/streptomycin, at 37°C in 5% CO_2_. For transient transfection of plasmids, cells were seeded at 3 x 10^5^ cells/well in 6-well plates and grown to 70–90% confluency. 2.5 µg of plasmid DNA were transfected per well using Lipofectamine 3000 (Invitrogen), following the manufacturer’s protocol. For transient siRNA transfections, cells were grown in 6-well plates to 60-80% confluency. 25 pmol of siRNA were transfected per well using Lipofectamine RNAiMAX (Invitrogen), following the manufacturer’s protocol. BLOCK-iT™ Alexa Fluor™ Red Fluorescent Control (Invitrogen) was used as a control and also co-transfected with RTN4 siRNA to identify transfected cells. The average transfection efficiency for each plasmid and siRNA was calculated from 3-5 independent experiments: 69.5% for GFP-NLS, 75.6% for Rtn4a-GFP, 62.2% for mCherry-REEP5, 72.4% for GFP-Rtn4b, 64.2% for BLOCK-iT Red Fluorescent Control alone, and 64.5% for the co-transfection of RTN4 siRNA with BLOCK-iT Red Fluorescent Control. To prepare cells for immunofluorescence analysis, transfected cells from each well were trypsinized using 450 µl of 1X Trypsin-EDTA solution (Sigma) at 24 h after transfection and transferred onto two acid-washed 18-mm square coverslips in 35-mm^2^ dishes with 2 ml of fresh culture medium. 12 h later, culture medium was removed and coverslips were processed for immunofluorescence. To prepare cell pellets for whole cell lysates, medium was removed from the 6-well plates at 24 h post transfection and replaced with 2ml/well of fresh culture medium. 12 h later, cells were trypsinized, spun down at 3500 rpm for 5 min, and washed twice in PBS.

### Immunofluorescence

36 h after transfection, coverslips were washed twice with phosphate-buffered saline (PBS), fixed with 4% paraformaldehyde for 15 min, and then subjected to three 5 min washes with PBS. Cells were permeabilized for 7 – 10 min with 0.25% Triton-X-100 and washed thrice with PBS for 5 min each. Blocking was performed for 1 hr at room temperature with 10% normal goat serum (Sigma) supplemented with 0.3 M glycine and then incubated overnight at 4°C with primary antibody diluted in 1.5% normal goat serum. Cells were washed thrice in PBS for 5 min each and incubated at room temperature for 1 hr with secondary antibodies diluted in 1.5% normal goat serum supplemented with 2.5 µg/ml Hoechst. Cells were then washed thrice in PBS for 5 min each followed by two brief washes with sterile water. Finally, coverslips were mounted in Vectashield (Vector Labratories) and sealed with nail polish. The following secondary antibodies were used at 1:500 dilutions: Alexa Fluor 488 and Alexa Fluor 568 conjugated Goat Anti-Rabbit IgG (H+L) and Anti-Mouse IgG (H+L) (Invitrogen) and Alexa Fluor 405 conjugated Goat Anti-Rabbit IgG (H&L) (Abcam 175652). The Triton-X-100 incubation step was skipped for experiments requiring non-permeabilized cells.

### Confocal Microscopy

Imaging was performed with a spinning-disk confocal microscope based on an Olympus IX71 microscope stand equipped with a five-line LMM5 laser launch (Spectral Applied Research) and switchable two-fiber output to allow imaging through either a Yokogawa CSU-X1 spinning-disk head or TIRF illuminator. Confocal images were acquired with an ORCA-Flash4.0 V2 Digital CMOS C11440-22CU camera (ImagEM, Hamamatsu) using an Olympus PlanAPO 100x/1.40 oil objective (for fixed cells) or Olympus UPlanFLN 60x/0.90 dry objective (for live cells). Z-axis focus was regulated using a piezo Pi-Foc (Physik Instrumente), and multiposition imaging was performed using a motorized Ludl stage. Image acquisition and all system components were controlled using MetaMorph software (Molecular Devices). All images were acquired using the same exposure time for a particular channel and experimental condition.

### Quantification from Fixed Cell Imaging

Unless otherwise noted, multiple z-stacks were acquired during fixed cell imaging for each cell, using a 0.2 µm z-slice thickness. Z-stacks were converted into maximum intensity projections in ImageJ, thresholded using MetaMorph software (Molecular Devices), and mean fluorescence intensity per cell was measured from thresholded images. Finally, mean fluorescence intensities from all cells were averaged for each condition. In order to quantify ER sheet and Golgi volumes, z-stacks were reconstructed and thresholded in 3D using MetaMorph software and the voxel volume was calculated based on the thresholded isosurface. Circularity/compactness of Golgi structures was measured from maximum intensity projections as previously described (Zahnleiter et al., 2015). Briefly, images were thresholded in MetaMorph and regions of interest were defined around Golgi clusters. The perimeter and area of every Golgi component were quantified and circularity of the Golgi apparatus was calculated using the formula 4π x [{sum(areas) ÷ sum(perimeters)}^2^]. Staining areas of Sec31A and COP-A were measured from thresholded images in MetaMorph. ERGIC53 staining distribution was quantified from maximum intensity projections in ImageJ by drawing three straight lines per cell (up to 15 µm in length) from the nuclear envelope to the cell periphery and measuring pixel intensities along the lines. Pixel intensities along the line scans were then averaged for all cells. For quantifying the colocalization of mCherry-LAMP1 (magenta) and Golgi (cyan), 3 - 4 single z-slices with prominent Golgi membranes were selected for both channels. Golgi images were thresholded in MetaMorph and total Golgi areas were quantified. Next, images from both channels corresponding to a particular z-plane were merged such that ‘white’ pixels corresponded to co-localization. The total area covered by white pixels was quantified manually in MetaMorph by drawing regions of interest around every white/merged component. To calculate the LAMP1 fraction that co-localized with the Golgi, the colocalized area (white) was divided by the total Golgi area for 3 – 4 selected z-planes per cell. These values were averaged from all z-planes for every cell to produce a mean value for the LAMP1 fraction that co-localized with the Golgi. For publication, images were cropped and pseudocolored using ImageJ, but were otherwise unaltered.

### Live Cell Imaging and Quantification

HeLa cells were transfected in 6-well plates as described previously. Cells from each well were trypsinized 24 h after transfection and divided into 8 wells of a chambered µ-Slide 8 Well (Ibidi – 80826) each containing 300 µl of fresh media. 12 h later, chambered slides were placed in a stage top incubator (Tokai Hit – INUBG2A-ZILCS) and confocal imaging was performed at a single focal plane. ER trapped LAMP1 was released by adding 40 µM D-Biotin to the media and time lapse imaging was started immediately. Images were acquired every 90 seconds and continued for 1 h 12 min. To quantify trafficking of LAMP1 through the secretory pathway, regions of interest of the same area were drawn around bright perinuclear LAMP1 puncta as well as dimmer perinuclear regions for every time point until 24 min post Biotin addition. Integrated densities of mCherry fluorescence were measured for the indicated regions in ImageJ. Integrated density for the brighter perinuclear LAMP1 signal was divided by the integrated density of the dimmer perinuclear LAMP1 signal at each time point to estimate accumulation of LAMP1 in the Golgi over time. Each of these ratios was then normalized to the lowest value in a time series for a given cell, averaged, and plotted as a function of time post ER release. Peaks were interpreted to represent the highest relative accumulation of LAMP1 in the Golgi for each condition.

### Western Blots

Whole-cell lysates were prepared from tissue culture cell pellets at 36 h post transfection using SDS-PAGE sample buffer supplemented with benzonase nuclease (Sigma, E1014) and boiled for 5 min. Proteins were separated on SDS-PAGE gels (4– 20% gradient) and transferred to PVDF membrane. Membranes were blocked in Odyssey PBS Blocking Buffer (Li-Cor, 927-40000). The primary and secondary antibodies were diluted in Odyssey PBS Blocking Buffer supplemented with 0.2% Tween-20. Anti-β-actin was used as a loading control. The secondary antibodies used were anti-mouse IRDye-680RD (Li-Cor 925-68070) and anti-rabbit IRDye-800CW (Li-Cor 925-32211) at 1:20,000. Blots were scanned on a Li-Cor Odyssey CLx instrument and band quantification was performed with ImageStudio. For a given sample, Rtn4a band intensity was normalized to the actin signal.

### EndoH Digestion

Whole cell lysate was prepared in RIPA buffer (50 mM Tris-HCl pH 7.5, 150 mM NaCl, 1% NP-40, 0.5% Sodium Deoxycholate, 0.1% SDS, 1 mM DTT, 1x Sigma Fast Protease Inhibitor cocktail) for EndoH digestion. Cell pellets were obtained at 36 h post transfection, resuspended in ice-cold RIPA buffer, and allowed to lyse for 30 min on ice. After lysis, cells were spun down at 14,000 rpm for 10 min at 4°C. The supernatant was stored at −80°C or used immediately. Total protein concentration was measured using the EZQ™ Protein Quantitation Kit (Invitrogen). 5-10 µg of total protein (9 µl of lysate) were treated with Endo H_f_ (New England Biolabs) for 1 h at 37°C, following the manufacturer’s protocol. Untreated controls were supplemented with GlycoBuffer3 (New England Biolabs). Reactions were stopped by treatment with SDS-PAGE sample buffer containing benzonase nuclease (Sigma, E1014) and boiling for 5 min. Samples were separated on SDS-PAGE gels (4–20% gradient) and transferred to PVDF membrane for immunoblotting.

### Concanavalin A Staining

Coverslips with cells were fixed 36 h post transfection and processed following the standard immunofluorescence protocol with or without permeabilization with 0.25% Triton-X-100. Blocking was performed with a 1% solution of glycoprotein free Bovine Serum Albumin (Sigma - A3059) in PBS for 1 h at room temperature. Coverslips were incubated with 10 µg/ml Biotinylated Concanavalin A (Vector Laboratories, a gift from Don Jarvis at the University of Wyoming) in PBS for 30 min at room temperature, followed by three 5 min washes with PBS. Cells were then stained with 5 µg/ml Texas Red Streptavidin (Vector Laboratories, a gift from Don Jarvis at the University of Wyoming) for 30 min at room temperature and washed thrice in PBS for 5 min each. Cells were stained for 5 min with Hoechst (5 µg/ml) and briefly washed thrice with PBS and twice with sterile water. Finally, coverslips were mounted in Vectashield (Vector Labratories) and sealed with nail polish.

### Primary Antibodies

| Antibody | Dilution |
| --- | --- |
| Anti-Rtn4a/Nogo A - Abcam (ab62024) | IF – 1:50; WB – 1:500 |
| Anti-Climp63/CKAP4 – Invitrogen (PA5-42926) | IF – 1:100 |
| Anti-Integrinβ1 – Invitrogen (PA5-29606) | IF – 1:100 |
| Anti-HLA-A – Santa Cruz Biotechnology (sc-390473) | IF – 1:60 |
| Anti-Integrinβ1 – Bethyl Laboratories (A303-735A) | WB – 1:1000 |
| Anti-HLA-A – Boster Bio (PB9376) | WB – 1:500 |
| Anti-GRASP65 – Invitrogen (PA3-910) | IF – 1:100 |
| Anti-Mannosidase II – EMD Millipore (AB3712) | IF – 1:200 |
| Anti-Sec31A – Santa Cruz Biotechnology (sc-376587) | IF – 1:60; WB – 1:250; IP – 20 μg/ml |
| Anti-VSV-G Tag – Invitrogen (PA1-29903) | IF – 1:250 |
| Anti-Calnexin – Enzo Life Sciences (ADI-SPA-860-D) | WB – 1:1000 |
| Anti-β-Actin – Abcam (ab8224) | WB – 1:1000 |
| Anti-ERp72 – ProSci (8211) | WB – 1:1000 |
| Anti-GRP78 - Assay Biotech (C0217) | WB – 1:500 |
| Anti-ERGIC53/LMAN1 – OriGene (TA502137) | IF – 1:100 |
| Anti-REEP5 – Abcam (ab76451) | IF – 1:100 |
| Anti-COP-A – Aviva Systems Biology (ARP51894_P050) | IF – 1:400 |
| Anti-Rtn4/Nogo – Santa Cruz Biotechnology (sc-271878) | IF – 1:75 |
| Anti-Rtn4 – Aviva Systems Biology (ARP46812_P050) | WB – 1:500 |

### Nucleofection and ELISA

Nucleofection of HeLa cells was performed using a Lonza 4D-Nucleofector™ device and Amaxa™ SE Cell Line 4D-Nucleofector™ X Kit (Lonza - V4XC-1024), according to the manufacturer’s protocol. 5.8 x 10^5^ cells were used per reaction. Post-nucleofection, cells were immediately seeded into 12 well plates containing 900 µl of medium per well. 12 h later, the conditioned medium was removed, spun down at 5000 rpm for 5 min at room temperature, and the supernatant was collected. The cells from each well were trypsinized, counted using a hemocytometer, and lysed in 50 µl of RIPA buffer. Both the media supernatant and whole cell lysate were subjected to sandwich ELISA. ELISAs were performed using FBLN5 (Fibulin-5) Human ELISA Kit from Fine Test (EH0772) and Thrombospondin 1 (TSP1) Human ELISA Kit from Invitrogen (BMS2100), following the manufacturer’s protocols. ELISA plates were read at 450 nm using a Wallac 1420 Victor2 Microplate Reader (Perkin Elmer, provided by Don Jarvis at the University of Wyoming). Whole cell lysate was diluted 1:10 in ELISA assay buffers. FBLN5 and TSP1 concentrations (ng/ml) were quantified in media (secreted) and whole cell lysate (intracellular) from standard curves. Total protein amounts (picograms) present in 900 µl of media and 50 µl of cell lysate were calculated. These values were normalized to the number of live cells per well. Secreted and intracellular protein amounts were added to calculate the total protein amount per cell. Data from three independent experiments were averaged. Average nucleofection efficiencies for GFP-NLS and Rtn4a-GFP were 71.6% and 82.2% respectively, calculated from three independent experiments.

### Immuno-isolation of Sec31A/COPII Vesicles

HeLa cells were nucleofected with plasmids expressing GFP-NLS or Rtn4a-GFP in triplicate (6.5 x 10^5^ cells per reaction) as previously described and seeded in T25 flasks containing 3 ml fresh culture medium. 24 h post-nucleofection, medium was removed and cells were scraped into 2.5 ml ice cold PBS (without Ca2+ and Mg2+) supplemented with 10 µg/ml each of leupeptin, pepstatin, and chymostatin. Cells were spun down at 2,000 rpm for 3 min at 4°C and the pellets were resuspended in 1.5 ml ice cold Homogenization Buffer (250 mM Sucrose, 25 mM KCl, 10 mM HEPES pH=7.2, 1 mM EGTA, and 1x Sigma Fast Protease Inhibitor cocktail), as previously described (Syed et al., 2017). Cell suspensions were homogenized by passing through a 25-gauge syringe needle (10 times) and homogenates were centrifuged (600 g for 5 min) twice at 4°C. The final supernatant (Post Nuclear Supernatant) containing intact membrane vesicles was used for immunoprecipitation using Pierce™ Protein A/G Magnetic Beads (Thermo Scientific), following the manufacturer’s protocol. A detergent-free wash buffer (Tris Buffered Saline) was used to maintain the integrity of immuno-isolated vesicles and all steps were performed at 4°C. 10 µg of anti-Sec31A antibody (Santa Cruz Biotechnology; sc-376587) or normal mouse IgG2a (Santa Cruz Biotechnology; sc-3878) were used per reaction. Proteins in immuno-isolated membrane vesicles were eluted in 50 µl of SDS-PAGE sample buffer and boiled for 5 min. 30 µl aliquots were separated on SDS-PAGE gels (4–20% gradient) and transferred onto PVDF membrane for immunoblotting with anti-Sec31A and anti-Rtn4a antibodies.

### Immuno Electron Microscopy

Cells from three independent transfection reactions were collected at 36 h post transfection and processed for immuno electron microscopy, following a previously described protocol (Phend et al., 1995). Briefly, cells were fixed with 4% paraformaldehyde + 0.5% glutaraldehyde in PBS at room temperature for 1.5 h and embedded in 2% low melting point agarose. Agarose cubes containing cells were rinsed with ddH2O three times for 5 min each and treated with 1% tannic acid in PBS for 1 h at room temperature. After three 5 min washes with ddH2O, agarose cubes were incubated with 2% uranyl acetate at room temperature for 1.5 h. Embedded cells were sequentially dehydrated in 10%, 20%, and 40% ethanol (10 min each), then 60% and 80% ethanol (20 min each), then 100% ethanol (three times, 15 min each). Next, infiltration was performed in 1:2 (v/v) LR White Resin (LRW):ethanol (100%) for 1.5 hr, 2:1 (v/v) LRW:ethanol (100%) for 1 hr, and then 100% LRW overnight at room temperature. Finally, cell blocks were sealed in beem capsules filled with LRW and polymerized overnight at 4–8°C under UV radiation. Polymerized blocks were trimmed and cut into 50 – 60 nm sections using an ultramicrotome and 3 sections were collected on each formvar-carbon coated nickel grid. For immuno-labeling, grids were blocked for nonspecific binding with 10% normal goat serum in PBS for 1 h at room temperature.

Grids were then incubated with rabbit anti-Rtn4a antibodies (Abcam) and mouse anti-Sec31A antibodies (Santa Cruz) diluted 1:20 in 1.5% goat serum overnight at 4°C, followed by three 5 min rinses in PBS. The grids were blocked again for 1 h at room temperature and incubated for 2 h with 1:20 Goat-Anti-Rabbit IgG antibodies (H+L) (EMS 25104) and 1:10 Goat-Anti-Mouse IgG (H+L) (EMS 25133) antibodies coupled with 6 nm and 15 nm gold particles, respectively, diluted in 1.5% goat serum. The grids were rinsed three times 5 min in PBS, stained with lead citrate for 30-60 sec, rinsed extensively in ddH_2_O, and air dried. All grids were imaged with a Hitachi H-7000 transmission electron microscope, equipped with a 4K×4K Gatan digital camera (Gatan, Inc., Pleasanton, CA). For each condition, 5 – 7 1513nm x 1513nm images were acquired from each section, for 7 - 10 sections from 3 grids. 8 – 10 sections were used from each polymerized block and 2 blocks were used per condition. All quantifications were performed in ImageJ and images were cropped to 500 nm x 500 nm for publication. All reagents for electron microscopy and secondary antibodies for immuno-gold labelling were purchased from Electron Microscopy Sciences.

### Statistics

Averaging and statistical analysis were performed for independently repeated experiments. Unpaired t-tests were performed using GraphPad Software to evaluate statistical significance. The p-values, number of independent experiments, and error bars are denoted in the figure legends.

## ACKNOWLEDGEMENTS

We thank David Fay and Amy Navratil for critical reading of the manuscript.

## AUTHOR CONTRIBUTIONS

Conceptualization, R.N.M., D.L.L.; Formal Analysis, R.N.M.; Investigation, R.N.M. performed all experiments; Z.Z. provided electron microscopy technical assistance; Resources, Z.Z.; Writing – Original Draft, R.N.M., D.L.L.; Writing – Review & Editing, R.N.M., Z.Z., D.L.L.; Funding Acquisition, D.L.L.; Supervision, D.L.L.

## Abbreviations

ER: endoplasmic reticulum
RER: rough ER
SER: smooth ER
tER: transitional ER
ERES: ER exit sites
Rtn: reticulon
EndoH: endoglycosidase H
ConA: concanavalin A
FBLN5: fibulin-5
TSP1: thrombospondin-1
ERGIC: ER-Golgi intermediate compartment
CNS: central nervous system
RHD: reticulon homology domain.

