## Supplemental Information for "Rtn4a promotes exocytosis in mammalian cells while ER morphology does not necessarily affect exocytosis and translation"

#### SUPPLEMENTAL FIGURE LEGENDS

**Figure S1. Rtn4a overexpression decreases ER sheet volume without causing ER stress (A-J) and reduces the levels of immature glycoproteins on the cell surface (K-N). Rtn4a overexpression increases cell surface localization, but not total levels, of Integrinβ1 and HLA-A in MRC-5 cells (O-V).** HeLa cells were transiently transfected with plasmids expressing GFP-NLS as a control or Rtn4a-GFP. **(A)** Whole cell lysates were immunoblotted for Rtn4a. It is typical for Rtn4a to run as a collection of bands ranging from 130 kDa to over 260 kDa (Li et al., 2004; Osborne et al., 2004). **(B)** Rtn4a band intensities were normalized to β-Actin loading controls. Averages from three independent experiments are shown. **(C)** Cells were immunostained for Rtn4a (red) and DNA (blue). **(D)** Rtn4a immunofluorescence intensity was quantified for 47-51 cells per condition. **(E-H)** Whole cell lysates were immunoblotted for ER stress-related chaperones calnexin, ERp72, and GRP78. Band intensities were normalized to β-Actin loading controls, and data from three independent experiments are shown. **(I)** Cells were stained for ER sheet marker CLIMP63 (red) and DNA (blue). **(J)** Quantitation of mean ER sheet volume based on CLIMP63 immunofluorescence, from 3D reconstructed confocal z-stacks. 54-57 cells were quantified per condition. **(K)** Non-permeabilized cells were stained for surface localized immature glycoproteins with concanavalin A (ConA). **(L)** ConA surface fluorescence staining intensity was quantified for 40 cells per condition. **(M)** Permeabilized cells were stained for total immature glycoproteins with ConA. **(N)** Total ConA staining intensity was quantified for 30-31 cells per condition. **(O-V)** MRC-5 cells were transiently transfected with plasmids expressing GFP-NLS as a control or Rtn4a-GFP. (**O-P**) Non-permeabilized MRC-5 cells were

stained for surface-localized Integrinβ1 (O, red) or HLA-A (P, red) and DNA (blue). (**Q**) Integrinβ1 surface fluorescence staining intensity was quantified for 25-31 cells per condition. (**R**) HLA-A surface fluorescence staining intensity was quantified for 20-25 cells per condition. (**S-T**) Permeabilized MRC-5 cells were stained for total Integrinβ1 (S, red) or HLA-A (T, red) and DNA (blue). (**U**) Total Integrinβ1 fluorescence staining intensity was quantified for 20-29 cells per condition. (**V**) Total HLA-A fluorescence staining intensity was quantified for 27-33 cells per condition.
Scale bars are 10 μm and images are maximum intensity projections of confocal z-stacks. Error bars represent standard deviation. **** p≤0.0001; ** p≤0.01; * p≤0.05; NS not significant.

**Figure S2. Rtn4 RNAi decreases Rtn4a and Rtn4b levels (A-C). REEP5 and Rtn4b overexpression decreases ER sheet volume without affecting cell surface localization of Integrinβ1 and HLA-A (D-Q). (A-C)** HeLa cells were transiently co-transfected with siRNA against Rtn4 and Block-iT fluorescent control siRNA or with Block-iT alone. **(A)** Whole cell lysates were immunoblotted for Rtn4. It is typical for Rtn4a to run as a collection of bands ranging from 130 kDa to over 260 kDa (Li et al., 2004; Osborne et al., 2004). **(B-C)** Rtn4a and Rtn4b band intensities were normalized to β-Actin loading controls. Averages from three independent experiments are shown. **(D-J)** HeLa cells were transiently transfected with plasmids expressing GFP-NLS (green) as a control or mCherry-REEP5 (red). **(D)** Cells were immunostained for REEP5 (yellow) and DNA (blue). **(E)** Average fluorescence intensity of REEP5 immunostaining was quantified for 20-27 cells per condition. **(F)** Cells were stained for ER sheet marker CLIMP63 (yellow) and DNA (blue). **(G)** Mean ER sheet volume based on CLIMP63 immunofluorescence was quantified from 3D reconstructed confocal z-stacks. 36-43 cells were quantified for each condition. **(H1-H2)** Non-permeabilized cells were stained for surface-localized Integrinβ1 (H1, yellow) or HLA-A (H2, yellow) and DNA (blue). **(I)** Integrinβ1 surface fluorescence staining intensity was quantified for 49-66 cells per condition. **(J)** HLA-A surface fluorescence staining intensity was quantified for 40-61 cells per condition. **(K-Q)** HeLa cells were transiently transfected with plasmids expressing GFP-NLS as a control or GFP-Rtn4b (green). **(K)** Whole cell lysates were immunoblotted for Rtn4. **(L)** Rtn4b band intensities were normalized to β-Actin loading controls. Averages from three independent experiments are shown. **(M)** Cells were immunostained for ER sheet marker CLIMP63 (red) and DNA (blue). **(N)** ER sheet volume was quantified from 3D reconstructed confocal z-stacks of CLIMP63-stained cells. 25-37 cells were quantified per condition. **(O1-O2)** Non-permeabilized cells were stained for surface-localized Integrinβ1 (O1, red) or HLA-A (O2, red) and DNA (blue). **(P)** Integrinβ1 surface fluorescence staining intensity was quantified for 30-33 cells per condition. **(Q)** HLA-A surface fluorescence staining intensity was quantified for 35-43 cells per condition.

Scale bars are 10 μm. Images are maximum intensity projections of confocal z-stacks. Error bars represent standard deviation. **** p≤0.0001; *** p≤0.001; ** p≤0.01; NS not significant.

**Figure S3. Rtn4a overexpression leads to more rapid trafficking of mCherry-LAMP1 into the Golgi and increases Golgi volume (A-F). REEP5 and Rtn4b overexpression lead to Golgi fragmentation, with REEP5 causing Golgi enlargement (G-I). (A-E)** HeLa cells were transiently co-transfected with plasmids expressing GFP-NLS as a control or Rtn4a-GFP (green) and the RUSH construct Str-Ii_LAMP1-SBP-mCherry (magenta). ER-trapped LAMP1 was released by addition of 40 μM D-Biotin to the growth media. **(A-B)** Cells were fixed 1-2 minutes after biotin addition and immunostained for Golgi marker GRASP65 (cyan). The LAMP1/Golgi merges in the yellow rectangles are shown magnified in (B). Images are representative single confocal z-planes. **(C-D)** Cells were fixed 5-6 minutes after biotin addition and immunostained for Golgi marker GRASP65 (cyan). The LAMP1/Golgi merges in the yellow rectangles are shown magnified in (D). Images are representative single confocal z-planes. **(E)** Colocalization of LAMP1 (magenta) and Golgi (cyan) generates white pixels in merged images. To calculate the Golgi fraction co-localized with LAMP1, the colocalized area (white) was divided by the total Golgi area (cyan) for 12-21 cells per condition. **(F)** HeLa cells were transiently transfected with plasmids expressing GFP-NLS as a control or Rtn4a-GFP (green) and stained for the Golgi marker GRASP65 (red). 3D reconstructed confocal z-stacks show Golgi enlargement in Rtn4a-transfected cells. **(G-I)** HeLa cells were transiently transfected with plasmids expressing GFP-NLS as a control (green), GFP-Rtn4b (green), or mCherry-REEP5 (red). **(G)** Cells were stained for Golgi marker GRASP65 (yellow) and DNA (blue). Images are maximum intensity projections of confocal z-stacks. **(H-I)** Golgi circularity (H) and volume (I) were quantified based on GRASP65 staining for 30-33 cells per condition. Volume was quantified from 3D reconstructed confocal z-stacks.

Scale bars are 10 μm. Error bars represent standard deviation. **** p≤0.0001; *** p≤0.001; * p≤0.05; NS not significant.

**Figure S4. Rtn4a overexpression disperses the perinuclear distribution of ERGIC53 and COP-A.** HeLa cells were transiently transfected with plasmids expressing GFP-NLS as a control or Rtn4a-GFP (green). **(A)** Cells were immunostained for the ER-Golgi intermediate compartment marker ERGIC53 (red), along with ER sheet marker CLIMP63 and DNA (blue). **(B-C)** To quantify the distribution of ERGIC53 staining, intensity values were measured along straight lines (up to 15 μm in length) drawn from the nuclear envelope to the cell periphery. In (B), representative cells and lines are shown for the control ERGIC53 image from (A). In (C), pixel intensities along ERGIC53 line scans were averaged for 25-28 cells per condition and 3 lines per cell. **(D)** Average fluorescence staining intensity of ERGIC53 was quantified for 31 cells per condition. **(E)** Cells were immunostained for COPI vesicle marker COP-A (red) and DNA (blue). **(F-G)** Quantitation of mean COP-A fluorescence intensity (F) and area covered by COP-A staining (G) for 33 cells per condition.

Scale bars are 10 μm. Images are maximum intensity projections of confocal z-stacks. Error bars represent standard deviation. **** p≤0.0001; NS not significant.

**Figure S5. Sec31A and Rtn4a are present in tubulovesicular structures with Rtn4a overexpression narrowing ER tubule width.** HeLa cells were transiently transfected with plasmids expressing GFP-NLS as a control or Rtn4a-GFP. Cells were processed for TEM, and immuno-gold labeling was performed to label Rtn4 with 6 nm gold particles and Sec31A with 15 nm gold particles. **(A-B)** The left panels show representative immuno-TEM images of tubulovesicular structures in which both Rtn4a

and Sec31A are present. The right panels show the same images overlaid with cartoon representations of Sec31A (blue circle), Rtn4a (yellow 'W'), and membrane (black dashed lines). **(C)** Widths of Rtn4a-labeled ER tubules were measured by quantifying the distances between two lipid bilayers for 107-125 randomly selected regions per condition. The average diameter of Sec31A-containing vesicles (~65 nm) was unchanged upon Rtn4a overexpression (data not shown).
7-10 50 nm ultra-thin sections were analyzed per condition. Scale bars are 100 nm. Error bars represent standard deviation. **** $p \leq 0.0001$.

#### MOVIE LEGENDS

**Video S1. Rtn4a overexpression accelerates trafficking of LAMP1 from the ER.** HeLa cells were transiently co-transfected with plasmids expressing GFP-NLS as a control or Rtn4a-GFP and the RUSH construct Str-Ii_LAMP1-SBP-mCherry (red). ER-trapped LAMP1 was released by addition of 40 μM D-Biotin to the growth media. Live cell imaging was performed at 90-second intervals, for 62 minutes after D-Biotin addition. Representative videos of the trafficking of mCherry-LAMP1 in control **(A)** and Rtn4a-transfected **(B)** cells are shown. Time-lapse images were cropped and compiled in ImageJ.

### Figure S1

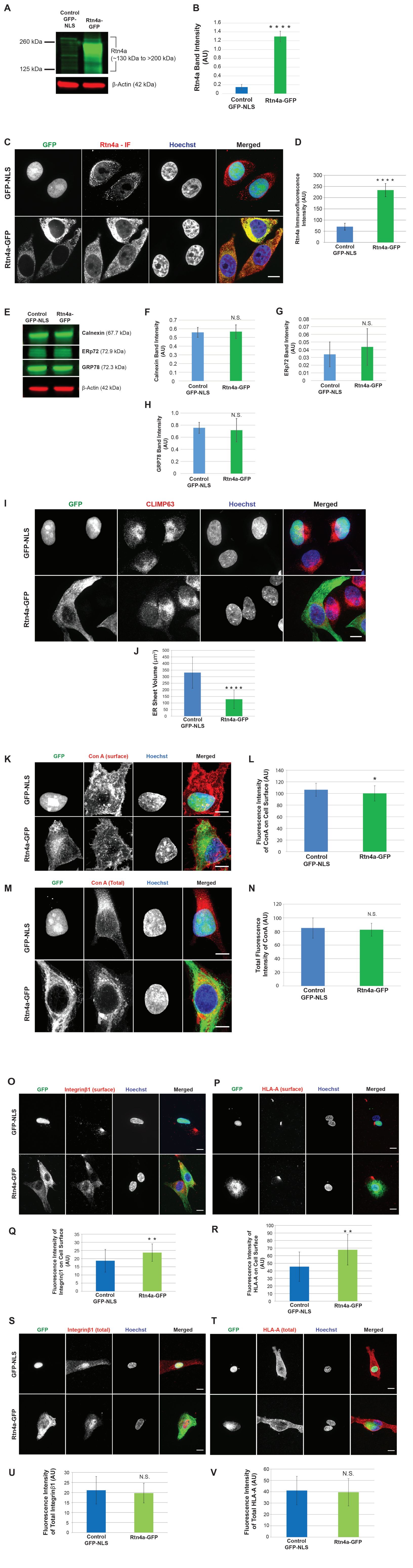

A
Control GFP-NLS
Rtn4a-GFP
260 kDa
125 kDa
Rtn4a (~130 kDa to >200 kDa)
β-Actin (42 kDa)
B
Rtn4a Band Intensity (AU)
Control GFP-NLS
Rtn4a-GFP
C
GFP
Rtn4a - IF
Hoechst
Merged
GFP-NLS
Rtn4a-GFP
D
Rtn4a Immunofluorescence Intensity (AU)
E
Calnexin (67.7 kDa)
ERp72 (72.9 kDa)
GRP78 (72.3 kDa)
β-Actin (42 kDa)
F
Calnexin Band Intensity (AU)
N.S.
G
ERp72 Band Intensity (AU)
H
GRP78 Band Intensity (AU)
I
CLIMP63
J
ER Sheet Volume (μm³)
K
Con A (surface)
L
Fluorescence Intensity of ConA on Cell Surface (AU)
M
Con A (Total)
N
Total Fluorescence Intensity of ConA (AU)
O
Integrinβ1 (surface)
P
HLA-A (surface)
Q
Fluorescence Intensity of Integrinβ1 on Cell Surface (AU)
R
Fluorescence Intensity of HLA-A on Cell Surface (AU)
S
Integrinβ1 (total)
T
HLA-A (total)
U
Fluorescence Intensity of Total Integrinβ1 (AU)
V
Fluorescence Intensity of Total HLA-A (AU)

### Figure S2

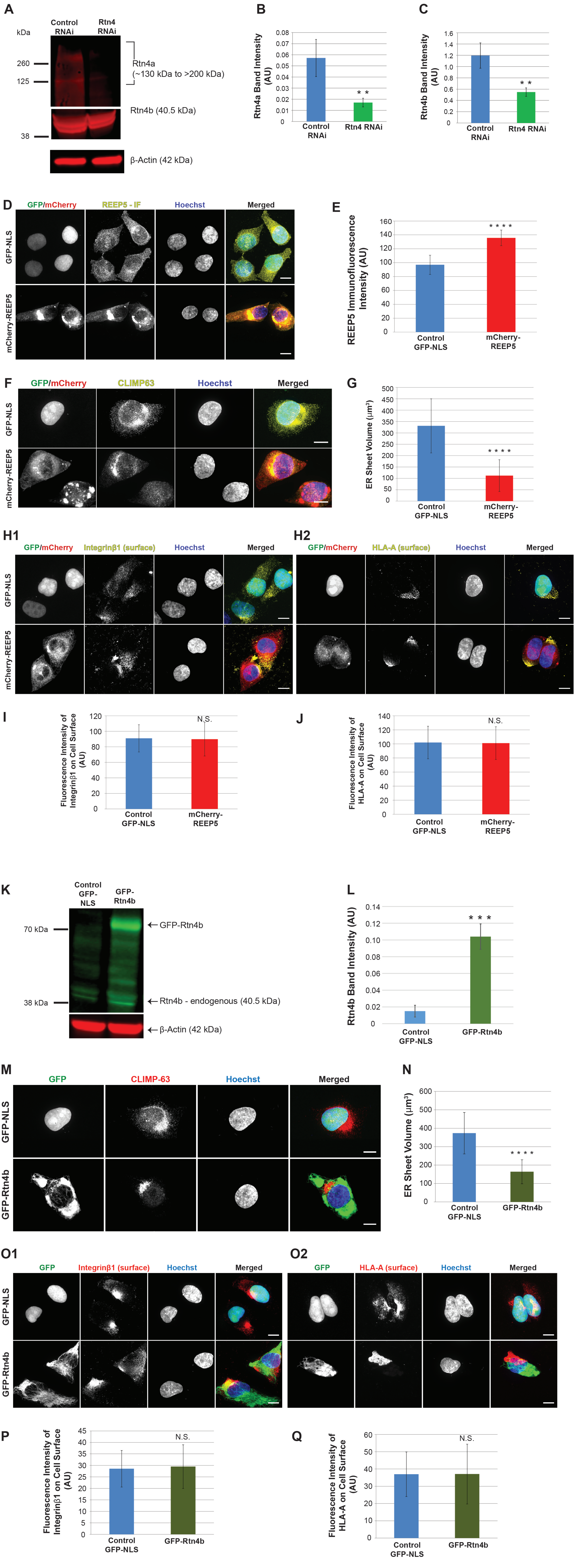

A
kDa
Control RNAi
Rtn4 RNAi
260
125
38
Rtn4a
(~130 kDa to >200 kDa)
Rtn4b (40.5 kDa)
β-Actin (42 kDa)
B
Rtn4a Band Intensity (AU)
Control RNAi
Rtn4 RNAi
C
Rtn4b Band Intensity (AU)
Control RNAi
Rtn4 RNAi
D
GFP/mCherry
REEP5 - IF
Hoechst
Merged
GFP-NLS
mCherry-REEP5
E
REEP5 Immunofluorescence Intensity (AU)
Control GFP-NLS
mCherry-REEP5
F
GFP/mCherry
CLIMP63
Hoechst
Merged
GFP-NLS
mCherry-REEP5
G
ER Sheet Volume (µm³)
Control GFP-NLS
mCherry-REEP5
H1
GFP/mCherry
Integrinβ1 (surface)
Hoechst
Merged
H2
GFP/mCherry
HLA-A (surface)
Hoechst
Merged
I
Fluorescence Intensity of Integrinβ1 on Cell Surface (AU)
N.S.
Control GFP-NLS
mCherry-REEP5
J
Fluorescence Intensity of HLA-A on Cell Surface (AU)
N.S.
Control GFP-NLS
mCherry-REEP5
K
Control GFP-NLS
GFP-Rtn4b
70 kDa
38 kDa
← GFP-Rtn4b
← Rtn4b - endogenous (40.5 kDa)
← β-Actin (42 kDa)
L
Rtn4b Band Intensity (AU)
Control GFP-NLS
GFP-Rtn4b
M
GFP
CLIMP-63
Hoechst
Merged
GFP-NLS
GFP-Rtn4b
N
ER Sheet Volume (µm³)
Control GFP-NLS
GFP-Rtn4b
O1
GFP
Integrinβ1 (surface)
Hoechst
Merged
O2
GFP
HLA-A (surface)
Hoechst
Merged
P
Fluorescence Intensity of Integrinβ1 on Cell Surface (AU)
N.S.
Control GFP-NLS
GFP-Rtn4b
Q
Fluorescence Intensity of HLA-A on Cell Surface (AU)
N.S.
Control GFP-NLS
GFP-Rtn4b

### Figure S3

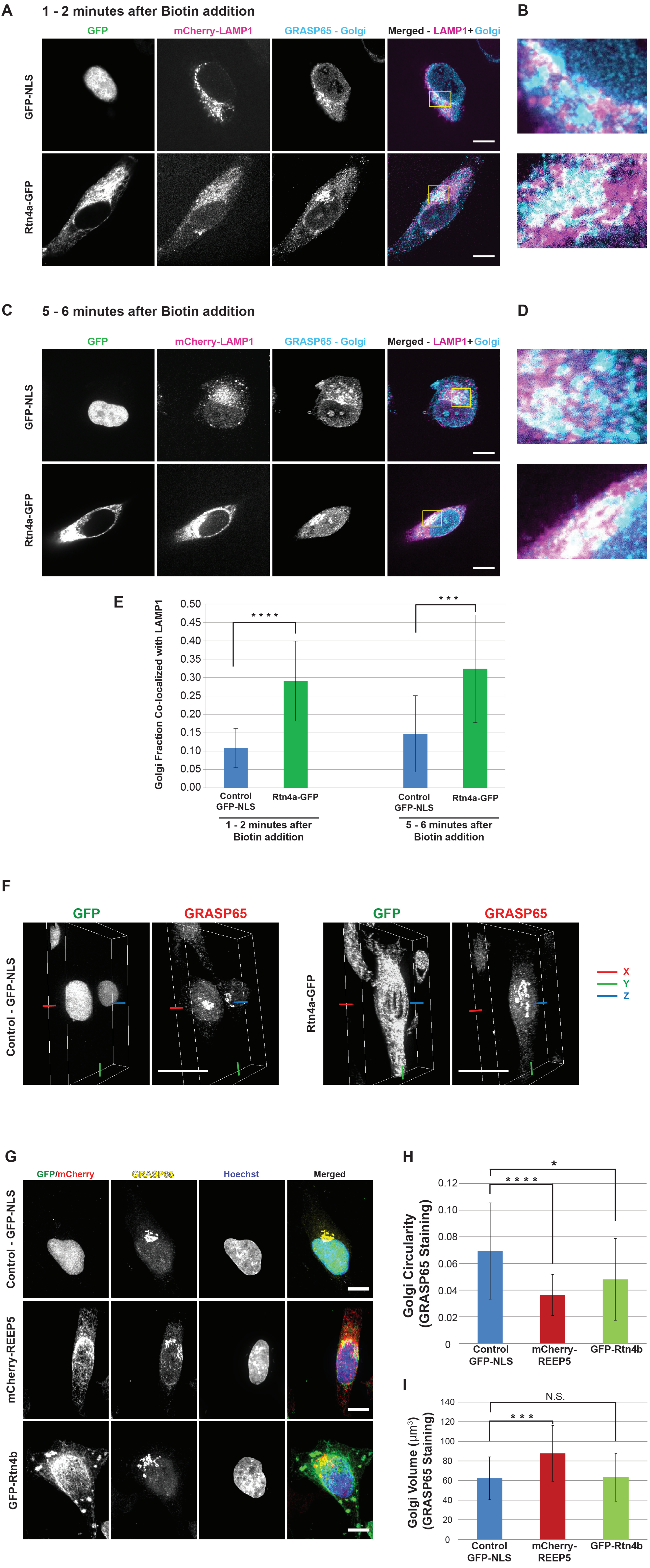

A
1 - 2 minutes after Biotin addition
B
GFP
mCherry-LAMP1
GRASP65 - Golgi
Merged - LAMP1+Golgi
GFP-NLS
Rtn4a-GFP
C
5 - 6 minutes after Biotin addition
D
E
Golgi Fraction Co-localized with LAMP1
0.50
0.45
0.40
0.35
0.30
0.25
0.20
0.15
0.10
0.05
0.00
* * * *
* * *
Control GFP-NLS
Rtn4a-GFP
1 - 2 minutes after Biotin addition
5 - 6 minutes after Biotin addition
F
GFP
GRASP65
Control - GFP-NLS
X
Y
Z
G
GFP/mCherry
GRASP65
Hoechst
Merged
mCherry-REEP5
GFP-Rtn4b
H
Golgi Circularity (GRASP65 Staining)
0.12
0.10
0.08
0.06
0.04
0.02
0
*
* * * *
Control GFP-NLS
mCherry-REEP5
GFP-Rtn4b
I
Golgi Volume (µm³) (GRASP65 Staining)
140
120
100
80
60
40
20
0
N.S.
* * *

### Figure S4

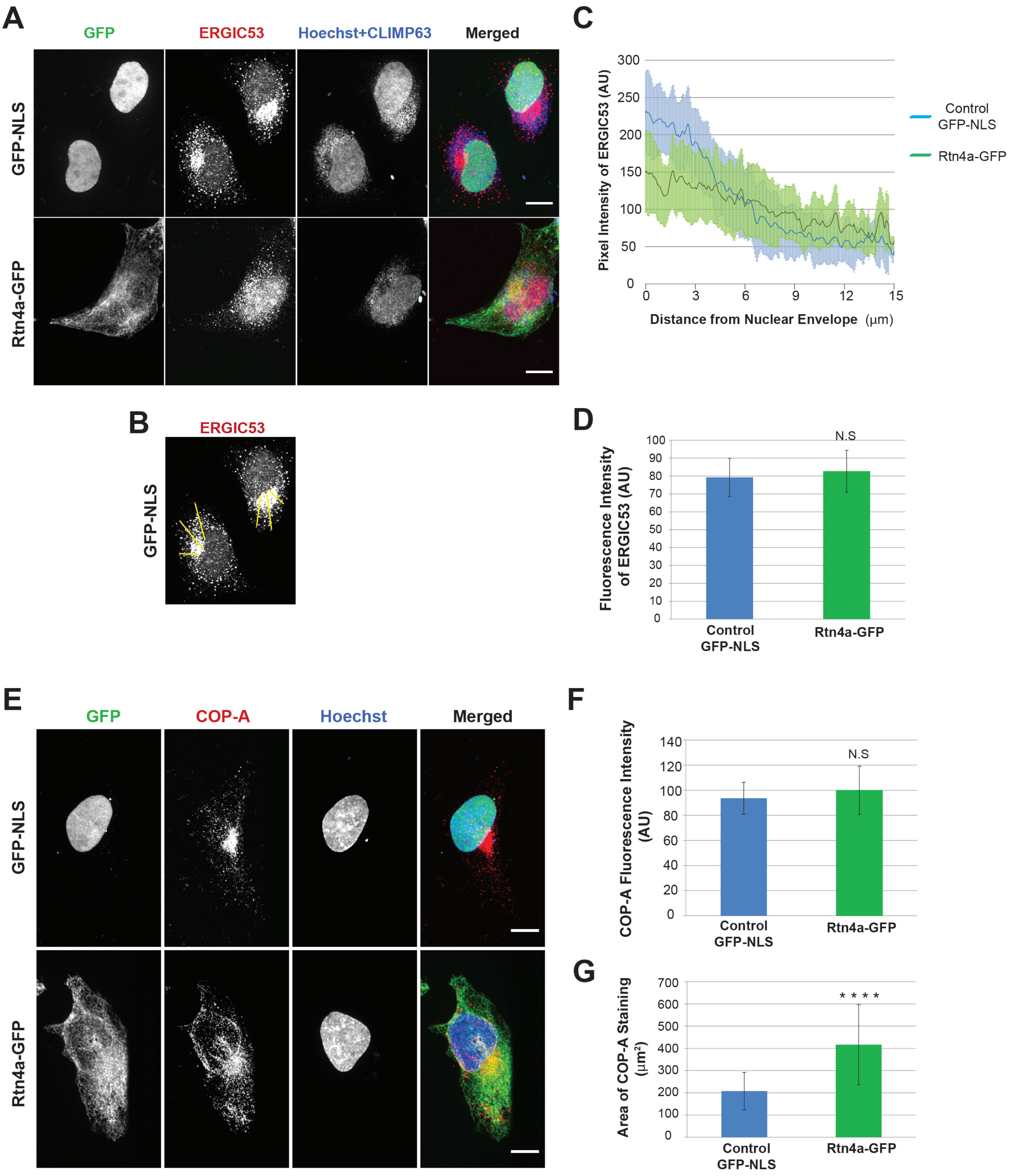
A
GFP
ERGIC53
Hoechst+CLIMP63
Merged
GFP-NLS
Rtn4a-GFP
B
ERGIC53
GFP-NLS
C
Pixel Intensity of ERGIC53 (AU)
300
250
200
150
100
50
0
0
3
6
9
12
15
Distance from Nuclear Envelope (μm)
Control GFP-NLS
Rtn4a-GFP
D
Fluorescence Intensity of ERGIC53 (AU)
100
90
80
70
60
50
40
30
20
10
0
N.S
Control GFP-NLS
Rtn4a-GFP
E
GFP
COP-A
Hoechst
Merged
GFP-NLS
Rtn4a-GFP
F
COP-A Fluorescence Intensity (AU)
140
120
100
80
60
40
20
0
N.S
Control GFP-NLS
Rtn4a-GFP
G
Area of COP-A Staining (μm²)
700
600
500
400
300
200
100
0
* * * *
Control GFP-NLS
Rtn4a-GFP

### Figure S5

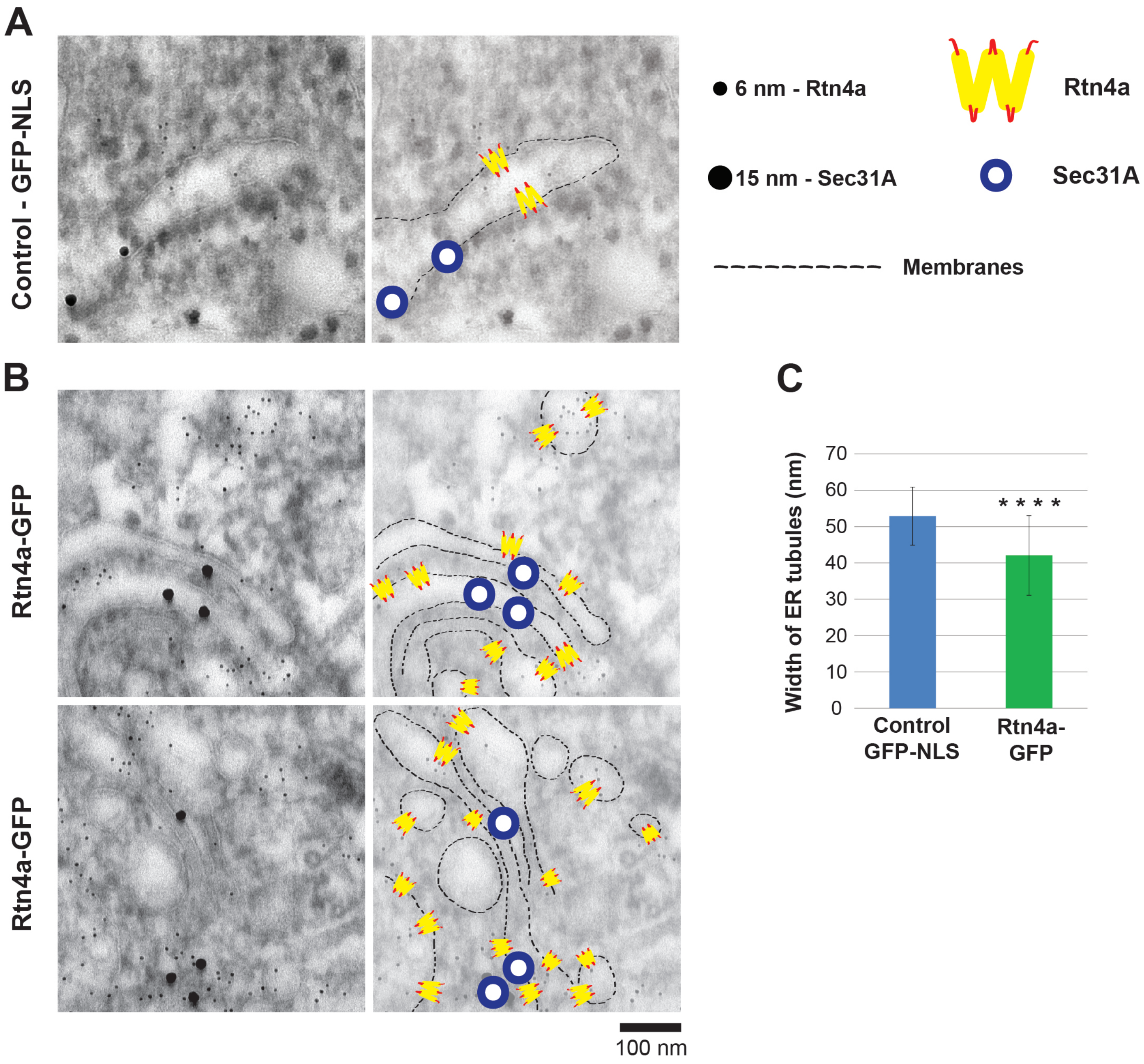
A
Control - GFP-NLS
6 nm - Rtn4a
15 nm - Sec31A
Rtn4a
Sec31A
Membranes
B
Rtn4a-GFP
Rtn4a-GFP
100 nm
C
Width of ER tubules (nm)
70
60
50
40
30
20
10
0
* * * *
Control GFP-NLS
Rtn4a-GFP
